# *Startle*: a star homoplasy approach for CRISPR-Cas9 lineage tracing

**DOI:** 10.1101/2022.12.18.520935

**Authors:** Palash Sashittal, Henri Schmidt, Michelle Chan, Benjamin J. Raphael

## Abstract

CRISPR-Cas9 based genome editing combined with single-cell sequencing enables the tracing of the history of cell divisions, or cellular lineage, in tissues and whole organisms. While standard phylogenetic approaches may be applied to reconstruct cellular lineage trees from this data, the unique features of the CRISPR-Cas9 editing process motivate the development of specialized models that describe the evolution of CRISPR-Cas9 induced mutations. Here, we introduce the *star homoplasy* model, a novel evolutionary model that constrains a phylogenetic character to mutate at most once along a lineage, capturing the *non-modifiability* property of CRISPR-Cas9 mutations. We derive a combinatorial characterization of star homoplasy phylogenies by identifying a relationship between the star homoplasy model and the binary perfect phylogeny model. We use this characterization to develop an algorithm, Startle (Star tree lineage estimator), that computes a maximum parsimony star homoplasy phylogeny. We demonstrate that Startle infers more accurate phylogenies on simulated CRISPR-based lineage tracing data compared to existing methods; particularly on data with high amounts of dropout and homoplasy. Startle also infers more parsimonious phylogenies with fewer metastatic migrations on a lineage tracing dataset from mouse metastatic lung adenocarcinoma.

**Code availability:** Software is available at https://github.com/raphael-group/startle

## 1 Introduction

Deriving the history of cell divisions that yield a multicellular organism from a single cell (*zygote*) is a key challenge in developmental biology. Such lineage tracing was accomplished for the 671 cells in *C.elegans* by J.E. Sulston in 1983 through elegant, but laborious experiments [Sul+83]. However, this direct experimental approach is intractable for more complex organisms such as mouse and human that contain trillions of cells. Several attempts have been made to infer the divisional history of cells, or a lineage tree, by measuring somatic mutations that accumulate in the genome of cells during cell division [Car+12; Beh+14; Lod+15; Bro+18; Tao+21]. However, somatic mutations typically accumulate gradually and sporadically across the genome, making it infeasible to infer high-resolution lineage tracing using such mutations as phylogenetic markers [MG19].

Recent developments in genome editing technologies, such as CRISPR-Cas9, have provided an alternative approach to lineage tracing. Specifically, technologies such as scGESTALT [McK+16; RGS18], ScarTrace [Ale+18], LINNAEUS [Spa+18], and others [Cha+19], allow researchers to induce heritable mutations at specific locations in the genome during development and at sufficiently high rate to enable high-resolution lineage tracing. While standard phylogenetic tree reconstruction algorithms such as such as neighbor joining [SN87], UPGMA [MS57], and hierarchical clustering have been used to infer single-cell phylogenies from lineage tracing data [Wag+18; Kal+18; Gon+22], several features of CRISPR-Cas9 lineage tracing data make the inference task quite difficult. First, CRISPR-Cas9 induced evolution has high rates of *homoplasy*, that is the same mutation occurs independently in multiple cells during the experiment [Kal+18; Spa+18]. Second, there is often a high rate of missing data (≈ 20-40%) and a large number of states per character/site (up to 50) in typical lineage tracing experiments [Raj+18; Jon+20; Cha+19]. Finally, the number of cells sequenced in these experiments can be upwards of several thousand, increasing the computational demands of tree reconstruction [Kon+22]. These challenges make reconstruction of single-cell phylogenies from current lineage tracing data a non-trivial computational task [MG19;WK20].

To address these challenges, several specialized methods have been developed to infer lineage trees lineage tracing data. For example, Cassiopeia [Jon+20] formulates phylogeny inference as a Steiner tree problem [ZK02] and presents three algorithms with varying scalability to solve the problem. Another method, LinTIMaT [ZLB20], uses a likelihood approach to integrate lineage information and gene expression from the same set of cells. GAPML [Fen+21] employs a statistical model specifically tailored for the scGESTALT lineage tracing technology. Several other phylogeny inference methods for lineage tracing data were recently benchmarked in the lineage reconstruction DREAM challenge [Gon+21]. One of the top performing methods from this challenge was Cassiopeia [Jon+20]; however, Cassiopeia uses heuristics to handle missing data which leads to poor performance with dropout rates that are typical in current lineage tracing technologies.

Here, we introduce a new evolutionary model, the *star homoplasy* model, that models the restrictions of the CRISPR-Cas9 mutational process. In particular, in the CRISPR-Cas9 system, the Cas9 protein is led by *guideRNAs* to create double-stranded breaks in specific sequences, known as *target-sites*, in the genome that are complementary to the guideRNA [AKL20]. This break is then repaired by the cell via error-prone DNA repair mechanisms that occasionally result in heritable insertions and deletions (indels) [PCL16]. Importantly, these induced mutations render the target-site distinct from the guideRNA, preventing further action of the Cas9 protein at that location in the genome. Thus, the defining characteristic feature of CRISPR-Cas9 induced evolution is that mutations are *non-modifiable*. That is, a target site (phylogenetic character) may mutate an arbitrary number of times during the course of evolution (homoplasy), but is constrained to mutate at most once along any lineage. To the best of our knowledge, there is no existing evolutionary model that explicitly models the non-modifiability property of CRISPR-Cas9 mutations, although this property is sometimes confused with irreversibility, a property of the Camin-Sokal evolutionary model [MG19;ZLB20]. An advantage of a model with restrictions on state transitions, like the star homoplasy model, is higher tolerance for missing data and errors since the allowed the state transitions impose constraints that facilitate imputation of missing data and error correction. For example, in cancer phylogeny, the infinite sites model [El-+15;JKB16;Pop+15;Jia+16;Des+15] or the Dollo model [El-18;Sat+20;Mal+19] have proved useful for helping address errors and missing data in bulk and single-cell DNA sequencing data.

We derive an algorithm, *Startle* (Star tree lineage estimator), to to infer a maximum-parsimony phylogenies from lineage tracing data under the *star homoplasy* model (Figure 1). *Startle* is based on a combinatorial characterization of star homoplasy phylogenies with a bounded number of homoplasies, a characterization derived from a relationship between the star homoplasy model and the two-state perfect phylogeny model. We show that *Startle* significantly outperforms state-of-the-art lineage tracing algorithms on simulations; with a particularly strong advantage on data with high amounts of dropout. On a recently published [Yan+22] lineage tracing data of metastatic lung adenocarcinoma in mouse models, we show that *Startle* recovers more parsimonious trees that supports fewer metastatic migrations compared to published results.

**Figure 1:**
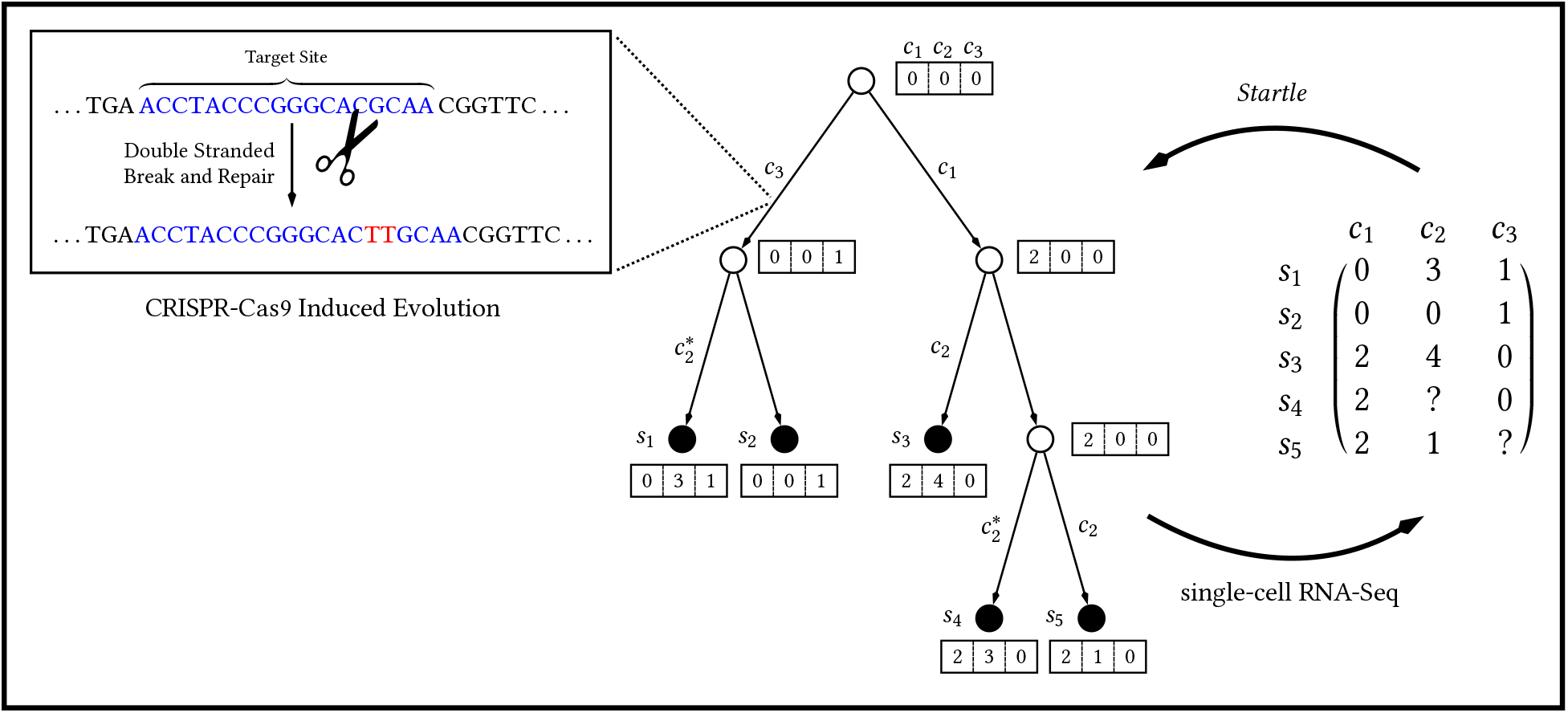
Overview of CRISPR-Cas9 lineage tree construction with *Startle*. (Left) CRISPR-Cas9 edits a genome at specific target sites, inducing heritable mutations in descendent cells. (Center) A cellular lineage tree. Labels on each vertex indicate the states of three characters (target sites) labeled *c*_1_, *c*_2_, and *c*_3_, and labels on each edge indicate character(s) that change state. Asterisks indicate homoplasies, or characters that change state multiple times on the tree. Character states for leaves of the tree, i.e. the extant cells, are measured via single-cell RNA-sequencing resulting in a character matrix (Right) indicating the state of each cell. ‘?’ indicate missing entries due to sequencing error. *Startle* imputes missing entries and reconstructs a lineage tree using the *star homoplasy* evolutionary model.

## 2 Methods

Suppose a lineage tracing experiment records mutations in *n* cells and *m* target-sites, or characters. We assume that each character begins in the unmutated 0 state and has *r* mutated states. We represent the measured mutations by a *n* × *m* character matrix *A* = [*a_ij_*], where entry *a_ij_* ∈ {−1, 0, …, *r*} is the state of character *j* in cell *i* with *a_ij_* = −1 indicating that the state of mutation *j* in cell *i* is missing, a common occurrence in lineage tracing data.

We represent the evolution of cells by a *phylogeny*, which is a rooted tree 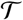 with *n* leaves, where the root 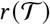 represents the founder cell in the experiment, each leaf in the leaf set 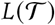 corresponds to an extant cell, and the internal vertices 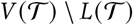 of 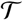 represent the ancestral cells. We define a *leaf labeling* 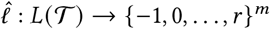 that labels a leaf 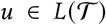 by the *i*^th^ row **a**_*i*_ of the character matrix *A*. Similarly, we define a vertex labeling 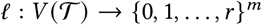 which represents the states of the characters in the cells associated with each vertex of 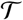, i.e. *ℓ*(*u*)_*j*_ is the state of character *j* in the cell associated with vertex 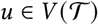. We say that vertex labeling *ℓ* is *consistent* with leaf labeling 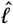 if 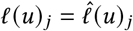 for all vertices *u* and characters *j* satisfying *ℓ*(*u*)_*j*_ ≠ −1. Importantly, a vertex labeling *ℓ* that is consistent with leaf labeling 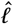 imputes the missing data of the character matrix *A* and infers the mutational state of unobserved ancestral cells in the phylogeny. We say that an edge 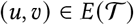 is a *mutation edge* with respect to vertex labeling *ℓ* if *u* and *υ* have a different state for some character *j* ∈[*m*], i.e. *ℓ*(*u*)_*j*_ ≠ *ℓ*(*υ*)_*j*_.

Given a character matrix *A*, our goal is to derive a phylogeny 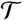 and vertex labeling *ℓ* that best describes *A* according to some scoring function, or evolutionary model, describing the scores of the state transitions on the mutation edges in 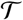. Below we derive a specialized evolutionary model, which we call *star homoplasy*, that models the unique characteristics of CRISPR-Cas9 induced evolution.

### 2.1 Star homoplasy model for CRISPR-Cas9 induced evolution

We introduce the *star homoplasy* evolutionary model that enforces the *non-modifiability* of mutations in CRISPR-Cas9 induced evolution. Specifically, each character in a *star homoplasy* phylogeny mutates at most once along a lineage. As such, mutation of a character is characterized by a transition from the unmutated state 0 to one of the mutated states. Multiple occurrences of the same mutation in a phylogeny is called *homoplasy* and, in the general case, the number of homoplasies for a mutation in the phylogeny is unbounded. A *bounded homoplasy* variant of our model is the *k-star homoplasy model*, in which each mutation can occur at most *k* times in the phylogeny.

To the best of our knowledge the star homoplasy model and its bounded homoplasy variant have not previously been described in the literature. To further describe the star homoplasy model and its relationship to existing models, we introduce *state-transition multigraphs*, a generalization of *state-transition graphs* (also known as character-state graphs) [SM92]. State-transition graphs are directed graphs in which the nodes are the states of the character and directed edges describe the permissible transitions between the states^1^. As an example, the most permissive model is the *finite states* model is which all transitions between all states of a character are allowed. Thus, the state-transition graph of the finite states model is the complete graph (Supplementary Figure S1c). Another example is the Camin-Sokal model that imposes an order on the mutated states and allows only those transitions that respect this order ([CS65], Chapter 10 [FF04]). Formally, the state-transition graph of the Camin-Sokal model is a tree in which all internal vertices except the root have exactly one incoming edge and one outgoing edge (Supplementary Figure S1b). The non-modifiability property of the star homoplasy model corresponds to a state-transition graph that is a *star tree* (Supplementary Figure S1c).

The star homoplasy model is closely related to the Camin-Sokal model. For binary (two state) characters, the star homoplasy and the Camin-Sokal models are equivalent. However, for characters with more than two states the two models are distinct as shown by the state-transition multigraphs for the two models (Supplementary Figure S1a vs. (Supplementary Figure S1b)). While several methods have been developed for the binary Camin-Sokal model [Fel78; Bon+17], the multi-state Camin-Sokal model is less studied; indeed popular phylogenetic packages such as PHYLIP [Fel93] implement only the binary Camin Sokal model. While multiple studies have used the binary Camin-Sokal model to infer trees from lineage tracing data [McK+16; Raj+18;RGS18], current lineage tracing technologies [Cha+19;Ale+18] yield characters with many states. We argue that the star homoplasy model – which enforces the non-modifiability property – is the appropriate model for phylogeny inference from such lineage tracing data than the multi-state Camin-Sokal model – which enforces only irreversibility.

Other evolutionary models impose restrictions on the number of homoplasies of a character in the phylogeny. The most common such restriction is the *no homoplasy* or *infinite sites* assumption. For a binary character, the infinite sites assumption says that the character may change at most once on the phylogenetic tree, while for more than two states, the assumption is that a character may change to a state at most once^2^. More generally, bounded homoplasy models impose an upper bound on the number of homoplasies. For example, the binary (or two-state) *k*-Dollo model bounds the number of transitions from state 0 to state 1 (gains) by one and the number of transitions from state 1 to state 0 (losses) by *k* [DJS86;El-18;Cic+20].

The *state-transition multigraph* (Figure 2) describes *both* restrictions on allowed transitions between the states of a character *and* restrictions on the number of transitions on the phylogeny. The state-transition multigraph is a directed multigraph where vertices are the states of a character, the directed edges indicate the allowed transitions between the states, and the multiplicity of each edge indicates the number of times that mutation is allowed to occur; i.e. the amount of allowed homoplasy. Edges with infinite multiplicity impose no restrictions on the number of transitions. Thus, the state-transition multigraph for the finite states model is a complete graph with all edges having infinite multiplicity (Figure 2g), while the state-transition multigraph for binary perfect phyologeny is a graph with a single directed edge (Figure 2a). The state-transition multigraph of the *star homoplasy* model is a star graph with edges that have infinite multiplicity (Figure 2e), while the *k-star homoplasy* model is represented by a star graph where each edge has multiplicity *k* (Figure 2d).

**Figure 2:**
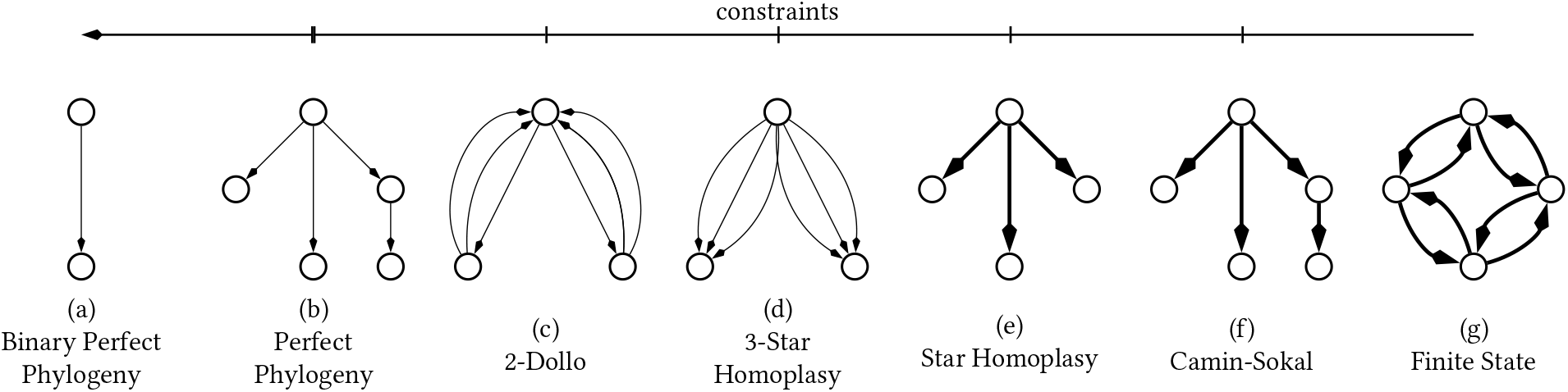
State-transition multigraphs for seven evolutionary models with (not-strictly) decreasing evolutionary constraints. The number of edges between two states denote the maximum number of times the transition is allowed to occur on the phylogeny; i.e. maximum number of homoplasies. Thick lines indicate edges with no restrictions on the number of times a transition can occur, i.e. infinite multiplicity.

### 2.2 Maximum parsimony under star homoplasy

We say that 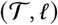 is a *k-star homoplasy phylogeny* provided: (1) every mutation edge of 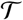 with respect to the labeling *ℓ* induces a state-transition that is allowed under the star homoplasy model; and (2) each state-transition occurs at most *k* times. Specifically, every mutation edge 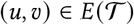 inducing a mutation in character *j* in *k*-star homoplasy phylogeny 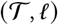 must have *ℓ*(*u*)_*j*_ = 0 and *ℓ*(*υ*)*_j_* = *s* for some nonzero state *s*. Moreover, a transition to state s for each character *j*, which we refer to as *mutation* (*j, s*), can occur at most *k* edges in the phylogeny.

We take the maximum parsimony approach [Far70] to evaluate a candidate *k*-star homoplasy phylogeny for a given character matrix *A*. Under the assumption that mutations occur independently along the branches of evolutionary tree, the cost, or score, 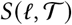 of a vertex labeling *ℓ* for a given phylogeny 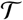 is

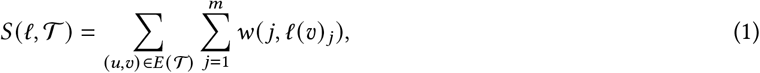

where *w*(*j, ℓ*(*υ*)_*j*_) is the weight of the transition from state 0 to *ℓ*(*υ*)*_j_* for character *j*. In the following, we use a weighted parsimony score with

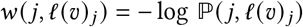

where 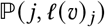 is the probability of a mutation. Setting *w*(*j, ℓ*(*υ*)_*j*_) = 1 corresponds to the standard (unweighted) parsimony.

We first pose the analog of the small parsimony problem [Fit71] of finding the optimal vertex labeling *ℓ* for a given phylogeny 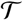 with a leaf labeling 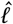.

#### Problem 1 (Small-Star Homoplasy Parsimony)

*Given a rooted tree* 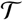, *a leaf labeling* 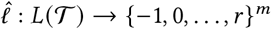 *and mutation weights w, find a vertex labeling* 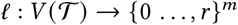 *such that ℓ consistent with* 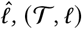 *is a k-star homoplasy phylogeny, and* 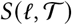 *is minimum overall such labelings*.

In general, the small parsimony problem for a given evolutionary model has multiple optimal solutions which can be enumerated using a dynamic programming algorithm called the Sankoff algorithm [SR75]. However, the specific constraints imposed by the *k*-star homoplasy model enforce a unique solution to the Small-SH Parsimony problem (Problem 1).

#### Theorem 1.

*Given a character matrix A with no missing data, a rooted tree* 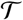, *and a leaf labeling* 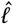, *there is a* unique *vertex labeling ℓ of minimum cost; i.e. the Small-SH Parsimony problem (Problem 1) has a unique solution*.

We prove the above theorem in Appendix A.4 with an algorithm that finds the unique solution to Problem 1 in 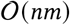 time.

Finally, we pose the analog of the large parsimony problem which is to find the most parsimonious *k*-star homoplasy phylogeny 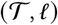 for an input matrix character matrix *A* (possibility with missing entries).

#### Problem 2 (Large-SH Parsimony)

*Given a character-matrix A and mutation weights w, find a k-star homoplasy phylogeny* 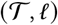, if one exists, that minimizes 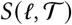.

We show that Problem 2 is NP-hard in Appendix B.2.

We say that a character-matrix *A* is a *k-star homoplasy matrix* if and only if there exists a *k*-star homoplasy phylogeny 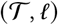 and a one-to-one mapping 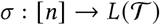 such that *ℓ*(*σ*(*i*))_*j*_ = *a*_*i, j*_ if *a_i, j_* ≠ −1 for all *i* ∈ [*n*]. Alternatively, we say that *A admits* a *k*-star homoplasy phylogeny. We conjecture that determining if *A* admits a *k*-star homoplasy phylogeny is NP-hard.

### 2.3 Combinatorial characterization of *k*-star homoplasy matrices

We derive a combinatorial characterization of *k*-star homoplasy matrices by drawing connections to binary perfect phylogenies. While this characterization does not yield a polynomial-time algorithm for the *k*-star homoplasy phylogeny, it is helpful for designing exact algorithms and heuristics. Specifically, we show that the problem of checking if a mutation matrix admits a *k*-star homoplasy phylogeny can be posed as a constrained variant of the binary perfect phylogeny problem with missing entries [PSS00].

We say that 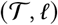 is a *binary perfect phylogeny* if every character has two states and every mutation occurs at most once. A binary perfect phylogeny matrix *A* is a binary matrix that admits a binary perfect phylogeny. We start with a well-known characterization of binary perfect phylogeny matrices [Gus14].

#### Theorem 2 ([Gus14])

*Suppose A* ∈ {0, 1}^*n*×*m*^ *is binary character-matrix and I*(*j*) ≔ {*i*: *a_i,j_* = 1, *i* ∈ [*n*]} *is the set of indices of the ‘1’ entries in the j^th^ column of A. A admits a perfect phylogeny if and only if for each pair j,j*′ *of characters, either: (i) I*(*j*) ⊆ *I*(*j*′); (*ii) I*(*j*′) ⊆ *I*(*j*)*; or (iii)* 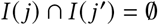.

We derive a connection between *k*-star homoplasy matrices *A* and binary perfect phylogeny by introducing new characters to describe the multiple occurrences (at most *k*) of each mutation (*j, s*) in the *k*-star homoplasy phylogeny. Specifically, given an *n* × *m* character matrix *A*, we define the *k-star binarization* of *A* to be an *n* × *mrk* binary matrix *B* = [*b*_*i*, (*j,s,p*)_] (where cell *j* ∈ {1,…, *m*}, state *s* ∈ {1, …, *r*} and occurrence *p* ∈{1,…, *k*}) in which each of the *m* multi-state characters in *A* is represented by *rk* binary characters. The binarization *B* is formally defined as follows.

#### Definition 1.

*A k-star binarization of a character matrix A* ∈ {−1, 0,…, *r*}^*n*×*m*^ *is a n* × *mrk binary matrix B* = [*b*_*i*, (*j,s,p*)_] *satisfying the following properties*.

1. *If a_i,j_* = 0 *for cell i and character j, then b*_*i*, (*j,s,p*_) = 0 *for all states s and all occurrences p*.
2. If *a_i,j_* = *s for cell i, character j and state s, then b*_*i*(*j,s, p*_) = 1 *for exactly one occurrence p and b*_*i*,(*j, s′, p*′_) = 0 *for all states s*′ ≠ *s*, *and occurrences p*′.
3. If *a_i,j_*, = −1 *for cell i and character j, then b*_*i* (*j,s,p*)_ = 1 *for at most one state-occurrence pair* (*s, p*).

Recall that the under the binary perfect phylogeny model all mutations are irreversible and, since there are only two states (0 indicates absence and 1 indicates presence of a mutation) for each character, the mutations are also non-modifiable. This leads to the following theorem.

#### Theorem 3.

*A character matrix A admits a k-star homoplasy phylogeny 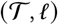 *with cost* 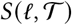 *if and only if there exists a k-star binarization B of A that admits a binary perfect phylogeny* 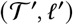 with the same cost*.

This theorem reduces the star homoplasy phylogeny problem to finding a *k*-star binarization of the character matrix with minimum cost. Once we have a *k*-star binarization *B* of a character matrix *A*, we can check if *B* admits a binary perfect phylogeny 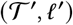, and construct the phylogeny, if it exists, in linear time [Gus91]. Further, as described in Section B.1, the binary perfect phylogeny 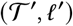 can be used to construct the *k*-star homoplasy phylogeny 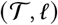 of character matrix *A* in linear time as well. However, finding a *k*-star binarization *B* of a character matrix *A* that admits a binary perfect phylogeny with minimum cost is NP-hard (Appendix B.2).

### 2.4 Startle

We describe two algorithms to solve the Large-SH Parsimony problem (Problem 2) under the star homoplasy model. The first algorithm, *Startle*-ILP, leverages the characterization described in Section 2.3 to formulate a mixed integer linear program (MILP) that solves Problem 2 exactly by finding the most parsimonious *k*-star binarization *B* of a character matrix *A* that admits a binary perfect phylogeny, if it exists. The second algorithm, *Startle*-NNI, employs tree operations to perform hill climbing [Rus10] in the space of star homoplasy phylogenies.

#### 2.4.1 Startle-ILP

We introduce binary variables *b_i,j,s,p_* for each cell *i*, character *j*, state s and occurrence *p* to model the entries of the *k*-star binarization matrix *B* of character matrix *A*. The MILP contains two types of constraints; (i) constraints to encode the three properties detailed in Definition 1 of the *k*-star binarization; and (ii) constraints to enforce that the *k*-star binarization *B* admits a binary perfect phylogeny. We describe constraints (i) and the objective function 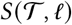 here. Appendix A.6 describes the constraints (ii).

##### Binarization Property 1

If *a_i,j_* = 0 for some cell *i*, character *j*, then *b*_*i*, (*j,s,p*)_ = 0 for all non-zero states *s* and occurrence *p*. We impose this property with the following constraints.

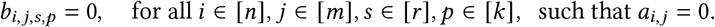

##### Binarization Property 2

If *a__i,j__* = *s* for some cell *i* and character *j* and state *s*, then *b*_*i*(*j,s,p*)_ = 1 for exactly one occurrence *p* and *b*_*i*(*j,s′,p*′)_ = 0 for all states *s*′ ≠ *s* and occurrences *p*′. We impose this property with following constraints for each cell *i*, character *j* and non-zero state *s* such that *a*_*i,j*_ = *s*.

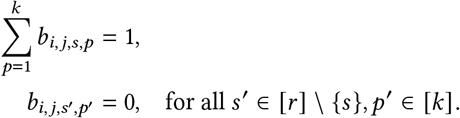

##### Binarization Property 3

If *a_i,j_* = −1 for some cell *i* and character *j*, then *b*_*i*,(*j,s,p*)_ = 1 for at most one state-occurrence pair (*s, p*). We impose this property with the following constraints for each cell *i* and mutation *j*.

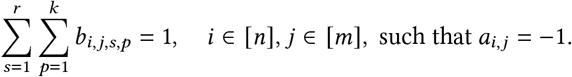

##### Objective function

Let 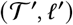 be the binary perfect phylogeny for the *k*-star binarization matrix *B*. Following Theorem 3, we need to minimize 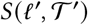 to solve Problem 2. Since a mutation can occur at most one in a binary perfect phylogeny, the cost of 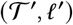 is the sum of the weight of all mutations that is present in at least one of the *n* cells. We introduce variables *x_j,s,p_* ∈ [0, 1] for each cell *j*, state *s* and occurrence *p* to indicate if column (*j, s,p*) in *B* has at least one nonzero entry. We model this as follows.

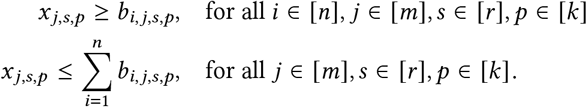

The objective of the MILP is set as follows.

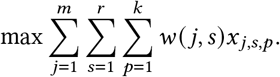

The above MILP has *O*(*nmrk*) binary variables, *O*(*nm*^2^*r*^2^*k*^2^) continuous variables and *O*(*nm*^2^*r*^2^*k*^2^) constraints. To improve the scalability of the MILP, we employ column generation and cut separation to introduce variables and constraints only as needed (details in Section A.6).

#### 2.4.2 *Startle*-NNI

We develop a second approach to solve the Large-SH Parsimony problem (Problem 2) that searches through tree space using sub-tree exchange operations. We start with a small set of candidate *binary* trees 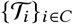 that we locally improve using sub-tree interchange operations. Specifically, we store a set of candidate trees and select one of the candidate trees uniformly at random. We then stochastically perturb the selected tree using random nearest neighbor interchange (NNI) operations [FF04]. With our randomly sampled and perturbed tree, we then apply a hill-climbing algorithm to minimize *S*(*ℓ, T*), the star homoplasy parsimony score, until we reach a local minimum. Once a local minimum is obtained, we check if the parsimony score improves on any of the candidate trees. If so, we update the candidate set by swapping our locally optimized tree with the candidate tree with the lowest parsimony score. Our approach resembles that taken in the popular phylogenetics package IQTree [Ngu+15].

Because of the non-modifiability property of the star homoplasy model, we can efficiently compute the star homoplasy parsimony score of all the trees in the NNI neighborhood of a particular tree during our hill climbing procedure. That is, since there are 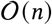 neighbors in the NNI neighborhood, a naïve approach that applies our linear time star homoplasy parsimony algorithm would take 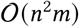 time in the worst case. In contrast, we devise an algorithm that takes 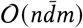 time to score all trees in the NNI neighborhood, where 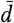 is the average depth of the current topology, by avoiding unnecessary re-computation (Theorem 5). We describe this and all other details of the algorithm in Appendix A.7.

## 3 Results

### 3.1 Simulated lineage tracing data

We compared *Startle*-ILP and *Startle*-NNI to neighbor joining [SN87] and Cassiopeia [Jon+20], which was one of the top performing methods in the DREAM challenge [Gon+21] for lineage tracing, on simulated data. We benchmark against all three modes of Cassiopeia [Jon+20]: the greedy algorithm (Cassiopeia-Greedy), the hybrid algorithm (Cassiopeia-Hybrid) and the ILP (Cassiopeia-ILP). We used the lineage tracing simulator in Cassiopeia to simulate data with *m* ∈ {10, 20, 30} target-sites (characters) and *n* ∈ {50, 100, 150, 200} cells. We varied the dropout probability *d* ∈ {0, 0.05, 0.15,0.2} which determines the amount of missing entries in the character matrix. Further details regarding the simulations are in Appendix A.1 and A.2. We compared the trees inferred by each method to the ground truth tree using three normalized metrics of tree dissimilarity: the Robinson-Folds (RF), Quartet, and Triple distances. We also evaluated the relationship between tree recovery accuracy and the weighted star homoplasy parsimony. Further details about these metrics and our comparison to ground truth are in Appendix A.3.

*Startle*-ILP and *Startle*-NNI substantially outperformed the other approaches on instances of moderate size with *n* = 150 or *n* = 200 cells and with realistic amounts of dropout *d* > 0 (Figure 3, Supplementary Figure S4). *Startle* found more accurate and parsimonious solutions than the Cassiopeia-Hybrid, Cassiopeia-Greedy, and neighbor joining algorithms. Neither Cassiopeia-ILP or Cassiopeia-Hybrid terminated after eight hours on the largest of these instances (Supplementary Table S1). Though we were able to run Cassiopeia-ILP and Cassiopeia-Hybrid on small instances of at most *n* = 100 cells and instances without dropout, their ability to scale is quite limited given that size of the ILP in these approaches scales exponentially in both the number of cells and characters. On the two medium sized instances where Cassiopeia-ILP did terminate, *Startle* was on average 30 times faster (Supplementary Table S1).

**Figure 3:**
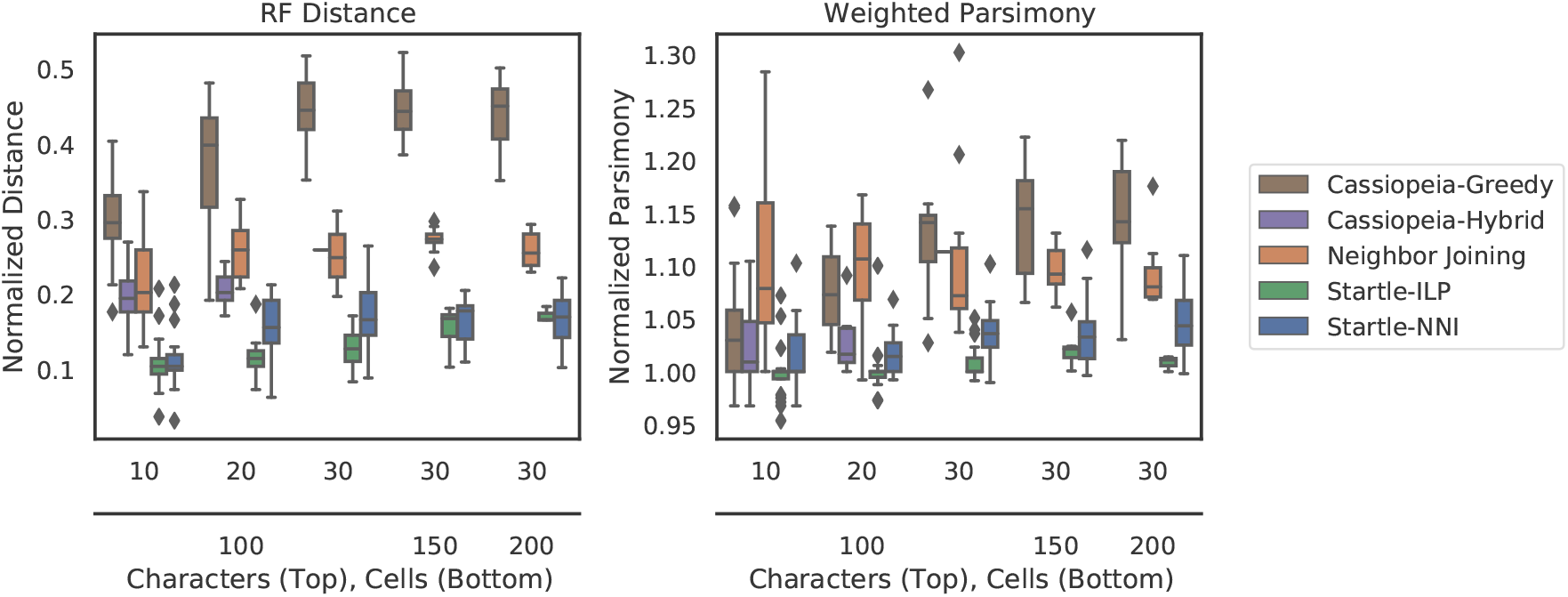
*Startle* outperforms existing methods on simulated lineage tracing data. (a) The normalized Robinson-Foulds (RF) distance between the tree inferred by each method and the simulated ground truth tree. (b) The weighted Star-homoplasy parsimony score for each method, normalized such that the simulated tree has score = 1. These simulated datasets were generated with mutation rate *p* = 0.1, dropout rate *d* = 0.2, and the indicated numbers of cells and characters.

On small instances with *n* = 50 or *n* = 100 cells, *m* = 10, 20, 30 characters and no dropout (*d* = 0), all methods, with the exception of Cassiopeia-Greedy, performed similarly and accurately recovered the ground truth trees. For example, both Startle-ILP and Cassiopeia-ILP recovered the ground truth *exactly* 22.5% of the time. On 36.1% of these instances, *Startle*-ILP and Cassiopeia-ILP inferred identical trees. However, with even a modest amount of dropout (*d* = 0.05), *Startle* substantially outperformed Cassiopeia-ILP, Cassiopeia-Hybrid, and neighbor joining, making *Startle* the top performing method by a large margin (Supplementary Figure S4b). On medium sized instances (*n* = 150, 200, and *m* = 30) and no dropout, all methods performed very similarly (Supplementary Figure S4a).

We also observed that a lower (weighted) star homoplasy parsimony score is correlated with improved tree recovery across all methods (Supplementary Table S2). Specifically, across all instances, the average Spearman’s rank correlation between median weighted parsimony score and the median RF distance is 0.84. This provides further evidence that the star homoplasy model is a useful model for these simulations since we can evaluate methods by how well they optimize star homoplasy parsimony score.

### 3.2 Lineage tracing in mouse metastatic lung adenocarcinoma

We apply *Startle* to analyze two datasets from a recently published CRISPR-based lineage tracing experiment in mouse models of metastatic lung adenocarcinoma [Yan+22]. The first dataset, `3513_NT_T1_Fam`, consists of 1227 total cells, including 969 cells from the primary tumor `3513_NT_T1`, 243 cells from a lymph node metastasis `3513_NT_N1` and 15 cells from a kidney metastasis `3513_NT_K1`. The published character matrix for this sample has 14 characters with a median of 3 states in each character. The second dataset, `3724_NT_T1_All`, consists of 21108 cells, including 14852 cells from the primary tumor `3724_NT_T1,` 3891 cells are from a soft tissue metastasis `3724_NT_S1`, 90 cells are from a liver metastasis `3724_NT_L1`, 1512 cells from a liver metastasis `3724_NT_L2` and 863 cells from a liver metastasis `3724_NT_L3`. The character matrix for this sample has 9 characters with a median of 38 states for each character. Due to the missing entries in this data (~6%), grouping cells that are indistinguishable by their measured mutations leads to significant reduction in the size of the problem, as described in Appendix A.5. We analyze `3513_NT_T1_Fam` using *Startle*-ILP and `3724_NT_T1_All` using *Startle*-NNI.

We find that *Startle* infers more parsimonious phylogenies than the phylogenies published in [Yan+22]. Specifically, *Startle* obtains score 315.51 for `3513_NT_T1_Fam` (Figure S5b) and 4715.5 for `3724_NT_T1_All` (Figure 4b) while the published phylogenies have scores 317.55 and 4827.43, respectively (Figure S5a, Figure 4a). Since the true phylogenies are unknown for this dataset, we evaluated the differences in the *Startle* phylogenies and the published phylogenies by examining the consistency between the phylogenies inferred by each method using subsets of cells. Specifically, we computed the normalized RF distance between the lineage tree inferred using only cells from the primary tumor and the lineage tree inferred using cells from all anatomical sites (both primary tumor and all metastases), but restricted to the cells of the primary tumor. We found that the phylogenies inferred by *Startle* are more consistent (normalized RF distance: 0.15 for `3513_NT_T1_Fam` and 0.525 for `3724_NT_T1_All`) than the published phylogenies (normalized RF distance: 0.22 and 0.6, respectively). This shows that the maximum parsimony star homoplasy model used in *Startle* yields a more robust phylogeny on this data.

**Figure 4:**
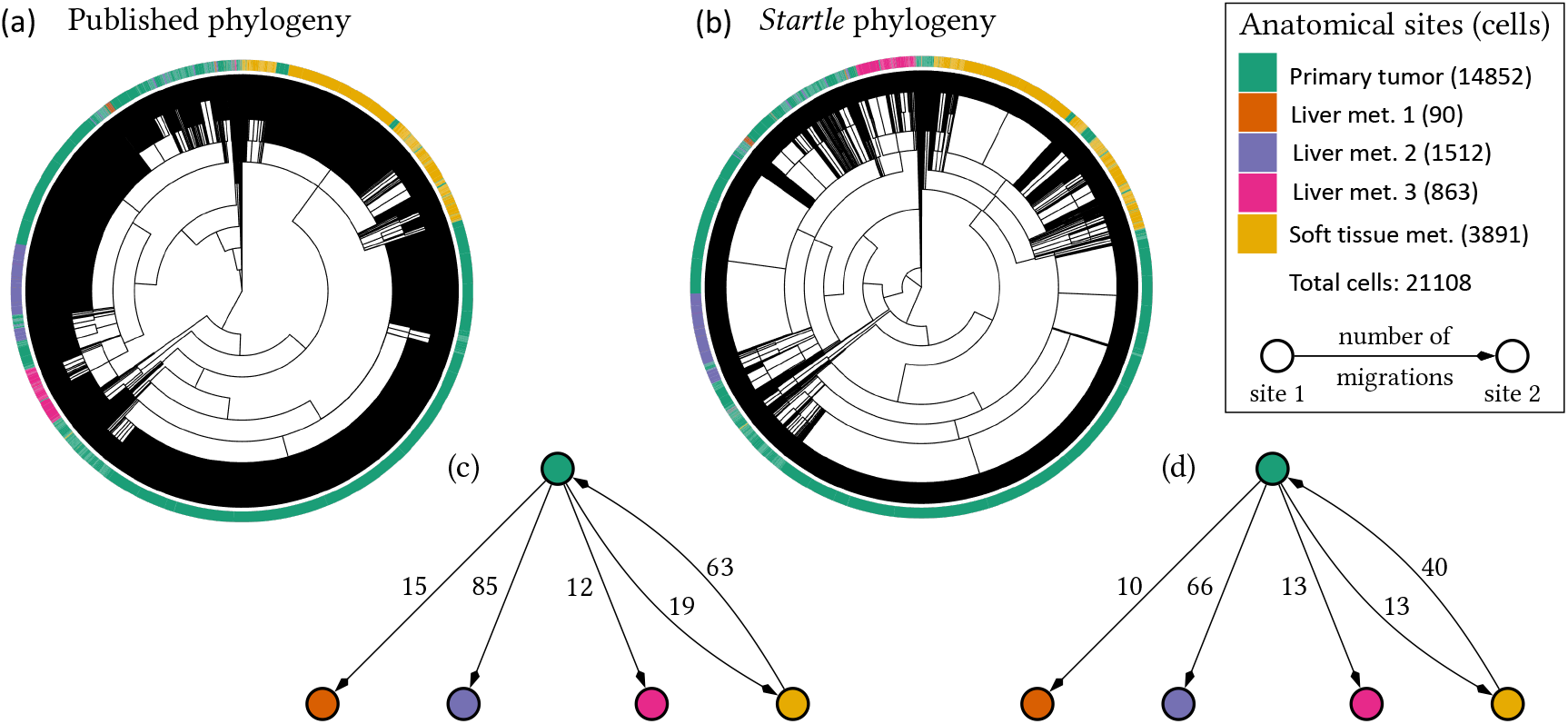
*Startle* infers more parsimonious lineage trees with fewer migrations in lineage tracing data from metastatic lung adenocarcinoma compared to the published phylogenies. (a) Published lineage tree and (b) *Startle* lineage tree for mouse lung adenocarcinoma sample `3724_NT_T1_All`. MACHINA migration graphs inferred using the (c) published lineage tree and (d) *Startle* lineage tree. Directed edges indicate the direction of migration and the edge weights denote the number of migrations between the anatomical sites.

The phylogenies inferred by *Startle* also yield a more parsimonious description of the metastatic process compared to the phylogenies from the original study. We calculated the minimum number of cellular migrations between anatomical sites using the maximum parsimony approach^3^ introduced in MACHINA [ESR18]. Specifically, given a *labeling ℓ_s_*(*u*) of the anatomic site for each leaf 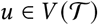 in the phylogeny 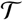, we find a label *ℓ_s_*(*u*) of anatomical site for each ancestral vertex 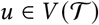 that minimizes the number of *migrations*, where a migration is an edge in the phylogeny that connects vertices labeled by distinct anatomical sites (details in Section A.8). The published phylogeny for `3724_NT_T1_All` suggests 213 migration while the *Startle* phylogeny supports a more parsimonious migration history with 156 migrations (Figure 4). Both the published and the *Startle* phylogeny agree that `3724_NT_T1_All` exhibits polyclonal seeding from the primary tumor `3724_NT_T1` to the four metastases (3724_NT_S1, 3724_NT_L1, 3724_NT_L2, 3724_NT_L3); i.e. multiple cells (or groups of cells) with different complements of mutations migrated from the primary tumor to these metastases. Both phylogenies also show reseeding from the soft tissue metastasis (`3724_NT_S1`) to the primary tumor (Figure 4). Similarly, the published phylogeny for `3513_NT_T1_Fam` results in 22 migrations from `3513_NT_T1` to `3513_NT_N1` and 2 migrations from `3513_NT_T1` to `3513_NT_K1`, while the *Startle* phylogeny supports fewer migrations (19 and 2, respectively, Figure S5). Both the published phylogeny and the *Startle* phylogeny agree that `3513_NT_T1_Fam` exhibits polyclonal seeding from the primary tumor `3513_NT_T1` to the two metastases, `3513_NT_N1` and `3513_NT_K1`.

## 4 Discussion

We introduced a novel evolutionary model, *star homoplasy*, for CRISPR-Cas9 based lineage tracing and an associated method *Startle* that reconstructs lineage trees by computing the maximum parsimony tree under this model. Star homoplasy models the observation that the CRISPR-Cas9 genome editing process induces at most one mutation at a target-site in a lineage, an assumption we term *non-modifiability*. We demonstrated that *Startle* recovers more parsimonious and accurate trees on simulated data with high rates of dropout and missing data, illustrating an advantage of the explicit constraints imposed by the star homoplasy model. *Startle* also inferred more plausible phylogenies on a recently published CRISPR-Cas9 based lineage tracing experiment from metastatic lung adenocarcinoma in mouse models [Yan+22].

There are multiple directions for future research. First, is to improve the scalability of *Startle* to thousands, or even millions of cells, while maintaining high accuracy. While *Startle*-ILP produces exact solutions, it does not scale to thousands of cells. Interestingly, during the present study we discovered that Cassiopeia-ILP exactly solves the weighted star homoplasy problem in the case when their potential graph is the boolean hypercube in *m* (the number of characters) dimensions [Jon+20]. This explains the strong performance of Cassiopeia-ILP on small instances, but since the boolean hypercube has 2^*m*^ vertices this approach is intractable for more than a few characters. *Startle*-NNI has fewer difficulties scaling, but could be further improved by adapting other tree rearrangement operations or other heuristics that have shown good performance in maximum parsimony and maximum likelihood software packages [Sta14;Gui+10]. Second, while *Startle* employs a maximum parsimony approach, we suspect that a maximum likelihood model for star homoplasy would have strong performance, particularly since maximum likelihood based methods outperform maximum parsimony methods in gene tree estimation [Fel78]. Third, the complexity of determining the existence of a *k*-star homoplasy phylogeny remains unknown; we conjecture this to be NP-hard. Finally, as lineage tracing technologies continue to develop, multiomic and spatial measurements of cell state along with lineage are being measured. Incorporating this auxiliary information into phylogenetic inference will create new opportunities and challenges. We argue that the star homoplasy model provides a framework for the development of improved algorithms for lineage tracing or other applications where the non-modifiablility assumption applies.

## A Supplementary methods

### A.1 Cassiopeia simulations

To construct our simulations, we first laid out a tree topology using Cassiopeia’s implementation of a forward birth-death process [Jon+20]. After laying out the tree topology, we used Cassiopeia’s Cas9-enabled lineage tracing data simulator to simulate the Cas9 recorder with *m* ∈ {10, 20, 30} target sites (characters) and *n* ∈ {50, 100, 150, 200} cells. We set an initial birth scale of 0.5, a fitness base of 1.3, and a stopping condition of *n* ∈ {50, 100, 150, 200} extant lineages. The birth and death waiting times were drawn from an exponential distribution. We allowed up to 25 states per character and drew mutation priors from an exponential distribution. We found that using a mutation rate *p* < 0.05 resulted in a nearly perfect phylogeny, which is trivial to infer with most computational methods. Conversely, a mutation rate *p* > 0.3 too quickly saturated the tree topology resulting in nearly all cells having the same barcode profiles. Thus, we set the mutation rate to *p* = 0.1 to achieve a balance between high and low saturation and so that there was a sufficient amount of homoplasy on these instances. For each parameter setting we constructed 20 instances using *s* ∈ {1,…, 20}as random seeds.

### A.2 Generating input for lineage tracing methods

*Startle*, Cassiopeia-Greedy, Cassiopeia-Hybrid and Cassiopeia-ILP take character matrices as input. In contrast, neighbor joining takes a *n* × *n* distance matrix *D* = [*d*_*i, i*′_] as input in which *d*_*i, i′*_ is an estimate of the evolutionary distance between cell *i* and *i*′ [SN87]. We set *d*_*i,i*′_ to be the normalized weighted Hamming distance implemented in Cassiopeia [Jon+20], defined as follows.

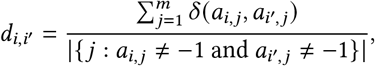

where

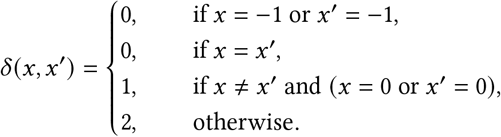

*Startle* additionally takes in input a value *k* for each mutation (character-state pair), indicating the maximum number of allowed occurrences for the mutation. We set the value of *k* for a mutation to the number of occurrences of that mutation in the (unweighted) Cassiopeia-Greedy solution. Cassiopeia-Greedy, Cassiopeia-Hybrid, and Cassiopeia-ILP were all ran with default parameters.

### A.3 Comparison to ground truth trees

To evaluate the quality of inferred trees against ground truth trees, we use three different tree dissimilarity metrics implemented by the TreeCmp tool [BGW12]. The input to our metrics is a ground truth tree 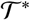 and an inferred tree 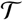. Both of these are specified in Newick format [CRV08].

To describe the metrics, we first define a function Contract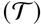 on edge-labeled phylogenetic trees that contracts all mutation-less edges. Specifically, for a mutation-less edge 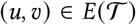, we contract the edge to single vertex *υ*′, and repeat this process until no mutation-less edges remain. This step is crucial for tree comparison metrics because because along a mutation-less edge the parent and child are indistinguishable. That is if (*u, υ*) is mutation-less, *u* and *υ* will be identical species when looking at the corresponding character matrix.

The Robinson-Foulds (RF) distance *d*_RF_(*T, T*\*) is a distance metric on trees defined on the set of induced bi-partitions in the input trees [RF81;BG11]. Specifically, with each edge 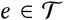 we associate a bi-partition 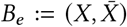 of its leaves using the equivalence relation *x* ~ *y* if *x* is connected to *y* in 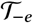, the forest resulting from the removal of edge *e*. We define the set 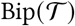 of bi-partitions for a tree 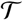 as 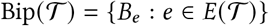. Then, the RF distance *d*_RF_(*T, T*\*) is defined as

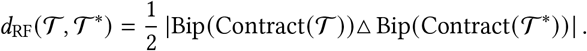

The Quartet distance 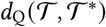 is computed by comparing the set of induced quartets on the input trees, when treating the input trees as unrooted [EMM85]. We define the set of quartets 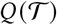 as the set of all 4-leaf sub-trees consistent with the unrooted topology of 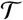. Then, the Quartet distance 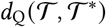 is defined similarly to the RF distance as

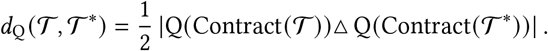

The Triples distance 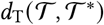 is computed by comparing the set of induced triples on the input trees [CPQ96]. We define the set of triples 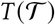 as the set of all rooted, 3-leaf sub-trees consistent with the topology of 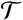. The Triples distance 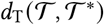 is then:

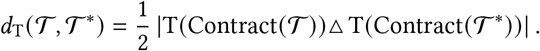

To allow for easy comparison of trees of different sizes, we use normalized variants of the tree distance metrics described in the previous sections. The normalized distance computation is implemented in the TreeCmp tool. For complete details on it, we refer the reader to their paper, but we will briefly summarize the normalized metrics here [BGW12]. For any distance metric 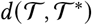 on trees each on *n*-taxa, we define a normalized distance as

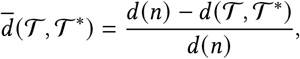

where *d*(*n*) is the expected distance between random trees using our metric *d*(·, ·). The Monte Carlo procedure to estimate *d*(*n*) is described in [BGW12].

### A.4 Small parsimony for star homoplasy

We efficiently solve the small parsimony problem under the star homoplasy model, Problem 1, using a dynamic programming algorithm similar to the Fitch and Sankoff algorithms [Fit71;SR75]. We note that any vertex labeling of individual characters, 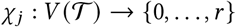 is easily extended to a vertex labeling of all characters 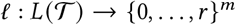 using the map

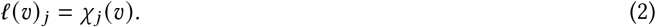

Further, we rewrite Equation 1 as

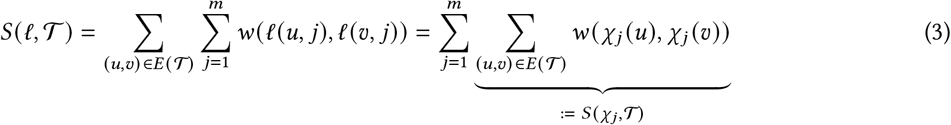

to observe that it suffices to minimize 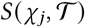 for each character separately.

We start by proving that Algorithm 1 minimizes 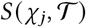 where the labeling *χ_j_* is consistent with a leaf labeling 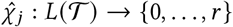 and is drawn from the set of admissible star homoplasy labelings of 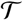.

#### Theorem 4.

*Algorithm 1 computes the the unique minimizer of* 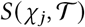 *under the star homoplasy model in time* 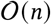.

*Proof*. Since we are performing a DFS, this algorithm takes linear time. It is also clear that the labeling of the leaves is unique since it must match 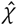. Theorem 4 then follows by induction on the size of the tree. Suppose *υ* is the root of 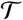 and let (*w*_1_, …, *w_k_*) be its children. By the induction hypothesis, *χ* is the unique, optimal labeling of the sub-tree rooted at *w_i_*. We then consider the size of the set *B* = {*χ*(*w_i_*): *χ* (*w_i_*) ≠ −1}. If *B* has size greater than or equal to 2, *χ*(*υ*) must be 0 as a non-zero state can’t mutate under the star homoplasy model. If the set *B* has size 1, all children *χ* (*w_i_*) = *c* for some *c* ∈ {0,1,…, *r*}. Then, *χ*(*υ*) must be either 0 or *c*, but it is only this latter labeling that minimizes 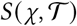. Finally, if the set *B* has size 0, all children of *υ* are labeled by missing entries; and again the most parsimonious solution is to label the root as missing as well. Since in each case, *χ*(*υ*) is uniquely determined up to a filling in of missing entries, we have proven our claim.

#### Algorithm 1

Small parsimony for star homoplasy.

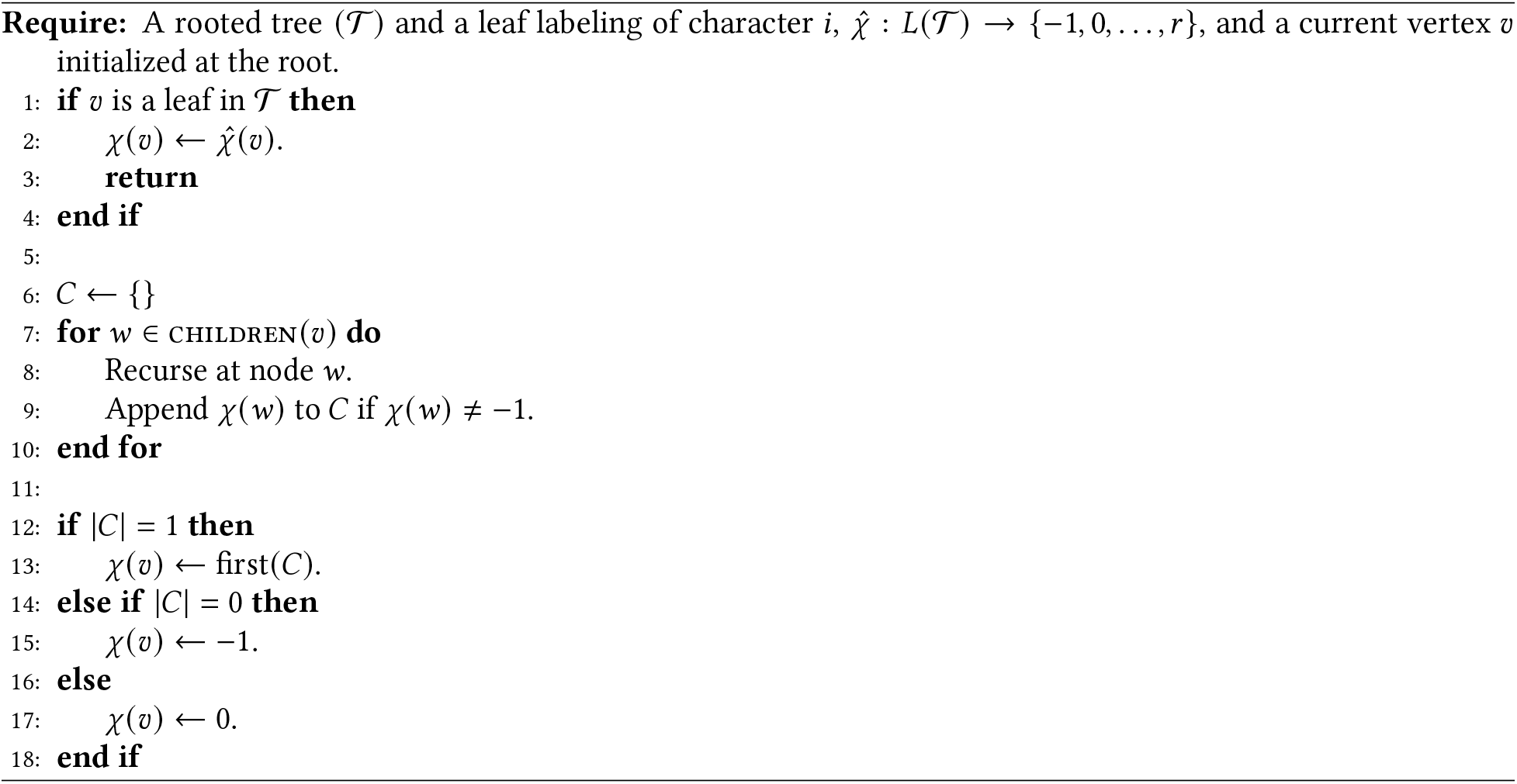

Since it suffices to minimizes 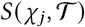 on each character separately, we simply run Algorithm 1 separately on each character and extend the vertex labeling using the map in Equation 2. This gives us the following corollary.

#### Corollary 1.

*Given a rooted tree* 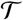 *and a leaf labeling* 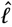, *we can find a vertex labeling ℓ that minimizes* 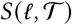 *such that* 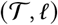 *is a star homoplasy phylogeny in* 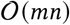 *time*.

Interestingly, the same algorithm finds a *k*-star homoplasy labeling. To see this, note that every mutation edge (*υ, w*) output by Algorithm 1 is forced – this follows from the case analysis in the proof of Theorem 4. Thus the number of mutations 0 → *s* in the vertex labeling by Algorithm 1 is a lower bound on the number of mutations 0 → *s* in *any* vertex labeling under the star homoplasy model. Thus it suffices to count the number of 0 → *s* mutations in the labeling output by Algorithm 1 and return HALT if any mutation occurs more than *k* times.

#### Corollary 2.

*Given a rooted tree* 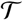 *and a leaf labeling* 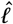, *there is an algorithm that finds a vertex labeling ℓ* minimizing 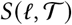 *such that* 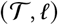 *is a k-star homoplasy phylogeny if one exists and otherwise outputs HALT in* 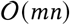 *time*.

### A.5 Reducing the size of the phylogeny inference problem

Here we describe a pre-processing step that reduces the size of the phylogeny inference problem without any performance degradation. Specifically, our pre-processing step identifies the smallest set of cells with distinguishable sets of mutations, removing the other cells, which can be placed on the tree afterwards. We first define a partial order relation ⪰ on the *n* cells based on their mutational profiles. Let *M*_1_(*i*) ={(*j, s*): *a_i, j_* = *s*} be the set of mutations (*j, s*) present in cell *i* and *M*_0_(*i*) ≔ {(*j, s*) : *a_i,j_* = *s*′, *s*′ ≠ *s*} be the set of mutations absent in cell *i*. We define a partial ordering on the cells as *i* ⪰ *i*′ if and only if *M*_1_(*i*) ⊇ *M*_1_(*i*′) and *M*_0_(*i*) ⊇ *M*_0_(*i*′). Said another way, we say that cell *i*′ precedes cell *i* if we can impute the missing entries of *i*′ to obtain *i*. We say that a cell *i* is *maximal* if there is no other cell *i*′ ≠ *i* such that *i*′ ⪰ *i*. Our pre-processing step identifies and outputs a set of maximal cells.

To construct our phylogeny, we solve the parsimonious star homoplasy problem on only the maximal cells. Then, we impute and place the remaining (not maximal) cells on the tree. Clearly this placement won’t increase the parsimony score of the phylogeny: every cell we place can be imputed to a cell already on the tree. As such, we can solve the maximum parsimony problem for all the *n* cells exactly while only considering the maximal cells during the computationally demanding phylogeny inference step. On real data from a recently published CRISPR-based lineage tracing experiment of metastatic lung adenocarcinoma in mouse models [Yan+22] (Section 3.2), we find 86 maximal cells (out of total 1227 cells) in `3513_NT_T1_Fam` and 1207 maximal cells (out of total 21108 cells) in `3724_NT_T1_All`, considerably reducing the computational cost of phylogeny inference.

### A.6 Integer linear program for inferring *k*-star homoplasy trees

We use the *set inclusion and disjointness* (SID) formulation stated in Theorem 2 to enforce that the k-star binarization *B* admits a perfect phylogeny [CRB10]. Recall that the 1-set of a column (*j, s, p*) is denoted by *I*((*j, s, p*)) = {*i*: *b*_*i*, (*j,s,p*) =1,*i*∈[*n*]_}. For all pairs of tuples (*j,s,p*), (*j*′, *s*′, *p*′), each representing a distinct column in the binary matrix *B*, we introduce two continuous variables: *y_j, s, p, j′, s′, p′_* and *z_j, s, p,j′,s′, p′_*. We force *y_j,s,p,j′, s′, p′_* = 0 if the 1-set *I*((*j, s, p*)) of (*j, s, p*) is not contained in the 1-set *I*((*j′, s′, p′*)) of (*j′, s′, p′*). Specifically, we enforce the following constraint.

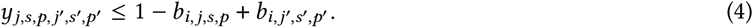

Along the same vein, we force *z_j, s, p, j′ s′, p′_* = 0 if the 1-set *I*((*j, s, p*)) of (*j, s, p*) and the 1-set *I*((*j, s, p′*)) of (*j′, s′, p′*) are not disjoint. We achieve this using the following constraint.

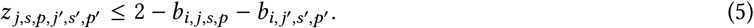

Finally, we enforce that every pair of columns ((*j, s, p*), (*j′, s′, p′*)) must be either related by containment or disjoint by imposing the following constraint.

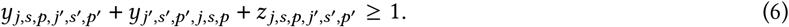

Algorithm 2 describes the column generation and cut separation steps to solve the MILP. Intuitively, we progressively introduce variables and constraints when we find violations of any of the above set inclusion and disjointness constraints until the ILP can be solved without any violations. Specifically, in the first iteration, we introduce variables *b_i,j,s,p_* but set the domains of *b_i,j,s,p_* ∈ {0, 1} if *p* = 1 and *b_i, j, s, p_* ∈ {0} if *p* > 1. At this point we do not impose any of the set inclusion and disjointness constraints (Eq.4, 5, 6). We solve the ILP and retrieve the *n* × *mrk* binary matrix *B*. We check for pair of columns (*j, s, p*), (*j′, s′, p′*) in *B* such that the 1-sets of *I*((*j, s, p*)) and *I*((*j′, s′, p′*)) are not disjoint or related by containment. If we do not find any such pairs, then we have solved the problem. For every pair of columns (*j, s, p*), (*j′, s′, p′*) in which we find a violation, we take the following steps.

a. Enforce *b_i,j,s,p_* ∈ {0, 1} and *b*_*i,j′,s′, p′*_ ∈ {0, 1} for all cells *i* ∈ [*n*].
b. Introduce variables *y_j,s,p,j′, s′, p′_, y_j′, s′, p′,j,s,p_*, *z_j,s,p,j′, s′, p′_*.
c. introduce constraints in Eq.4, Eq.5 and Eq.6.

#### Algorithm 2 Column generation and cut separation.

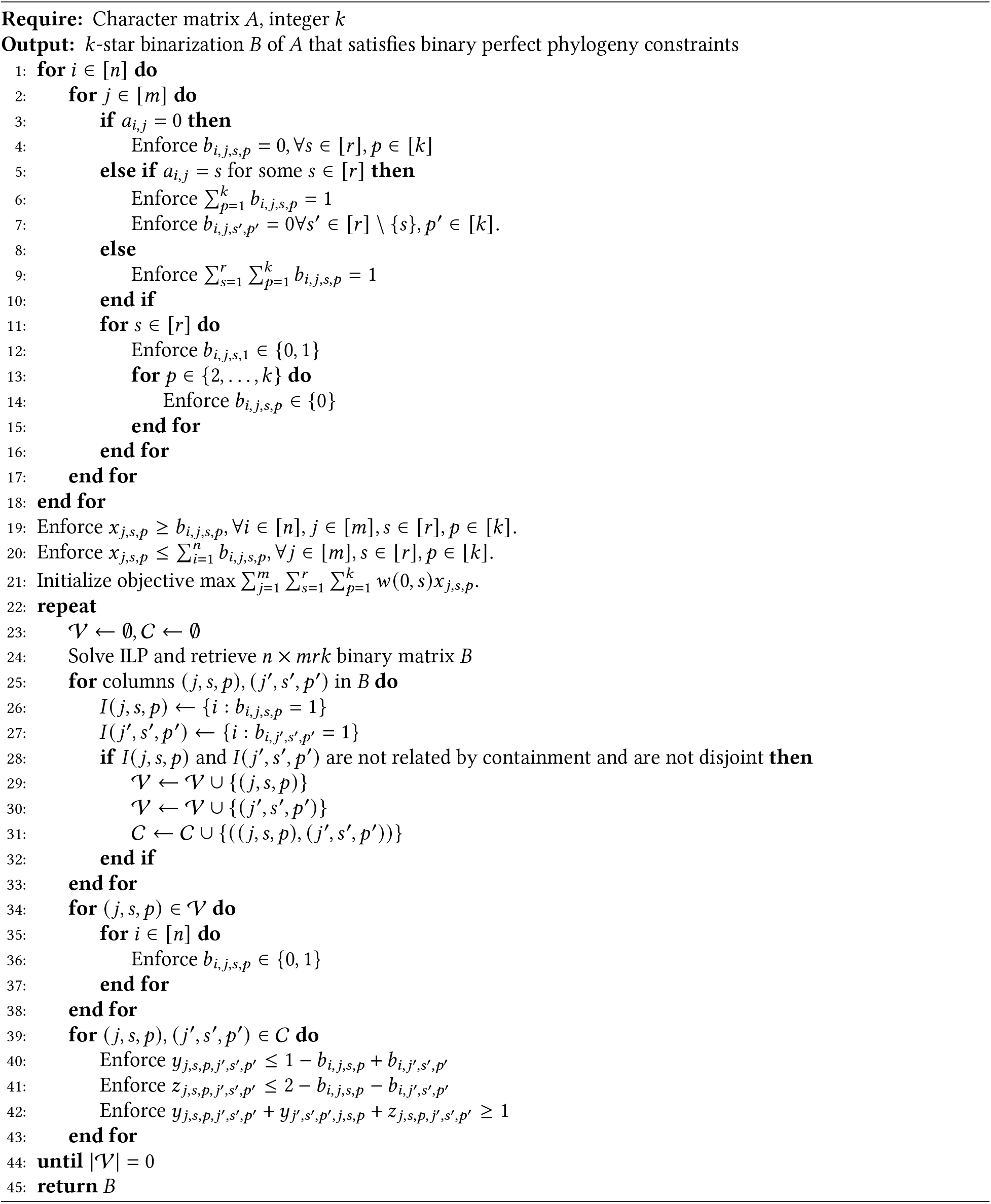

We then solve the new ILP again and repeat this process until no violations are found.

### A.7 Stochastic hill climbing with NNI moves

As described in our main text, our method maintains and improves a small set of candidate trees until no more improvements can be made. Specifically at every iteration, our algorithm randomly samples and stochastically perturbs a candidate tree on which it performs local optimization using a hill-climbing algorithm. This process is repeated until we do not see any improvement for a large number of iterations, *I*, which we set to 250 for our instances. This high-level approach is described in Algorithm 3.

#### Algorithm 3 Stochastic hill climbing for large parsimony under star homoplasy.

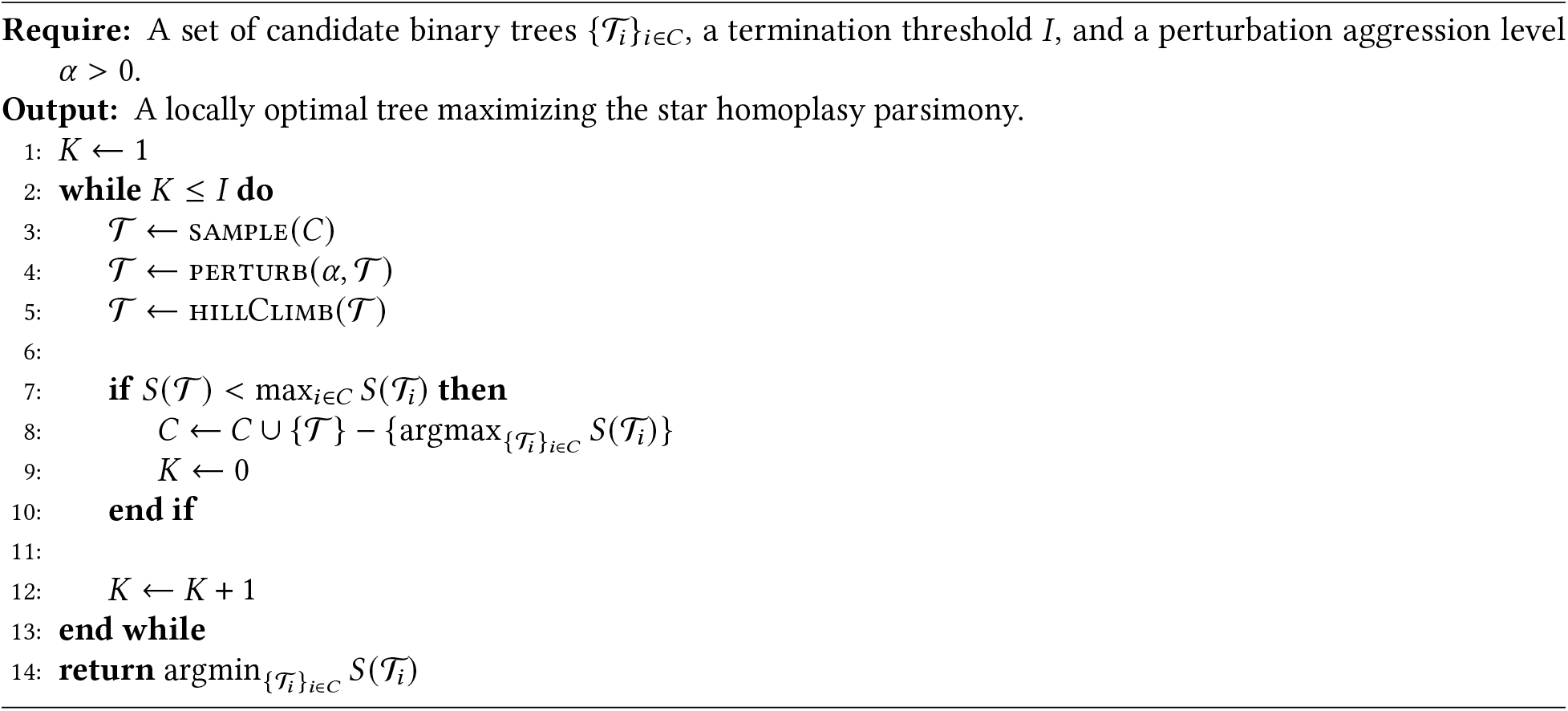

Let 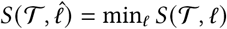 be the maximum score for any star homoplasy labeling of *l* consistent with our leaf set. As mentioned in Appendix A.4, this can be efficiently computed in linear time, though interestingly, it can be computed for all trees in the NNI neighborhood in 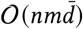 time, where 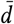 is the average depth of every leaf in the tree.

#### Theorem 5.

*We can compute* 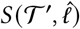 *for all trees* 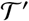 *in the NNI neighborhood of* 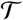 in 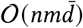 *time, where* 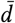 *is the average depth of a node in* 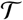.

*Proof*. To see this, suppose we have the optimal labeling for 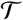 and wish to compute the optimal labeling for 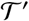 obtained by performing an NNI operation on 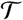 that swaps the sub-trees 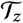 and 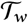 over edge (*u, υ*). For any nodes in the sub-trees of 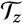 and 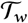, we do not need to recompute their optimal labeling since the sub-trees have not changed. Consider the path *P* from the root to vertex *υ*. By the previous observation, we do not need to recompute the optimal labeling of the trees rooted at children of *u* or *υ*. Further, the optimal labeling of any sub-trees off the path *P* do not need to be recomputed. Thus in total, we have to recompute the optimal labeling only for the nodes on *P*, which has size *d*(*υ*), the depth of node *υ*.

Since we can obtain the optimal labeling for 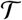 in 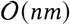 time, the total time taken is at most

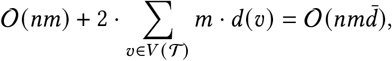

which proves the claim.

The hillClimb procedure computes 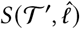 for all 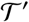 in the NNI neighborhood of 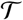 and selects the tree with the largest improvement. It continues until no more improvement can be made. The perturb procedure is implemented by performing selecting and executing a NNI uniformly at random *α* * 2(*n* – 3) times, where *α* > 0 is an aggression level hyper-parameter.

### A.8 Migration history inference

Here, we provide details of the algorithm to find the most parsimonious migration history for a given tumor phylogeny with metastatic samples. Let Σ be the set of all anatomical sites labeling and 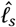 provide the anatomic site of origin of the cell associated with each leaf 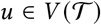 in the phylogeny 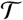. Let 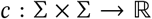 give the cost of migration between the anatomical sites. In the simplest case, we set *c*(*s, t*) = 1 if *s* ≠ *t* and *c*(*s, t*) = 0 if *s* = *t*. Let *f*(*υ, t*) be the minimum number of migrations in the subtree 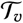 of 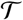 rooted at 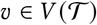 that can be attained when the site labeling of vertex *υ* is *t* ∈ Σ, i.e. *ℓ_s_*(*υ*) = *t*. *f*(*υ, t*) satisfies the following recurrence.

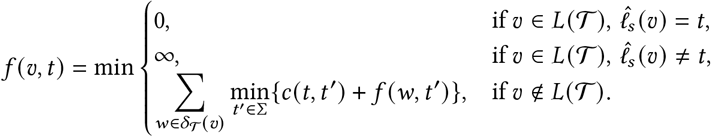

Algorithm 4 fills the cost matrix *f* using dynamic programming.

#### Algorithm 4 DpLabelingCost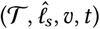

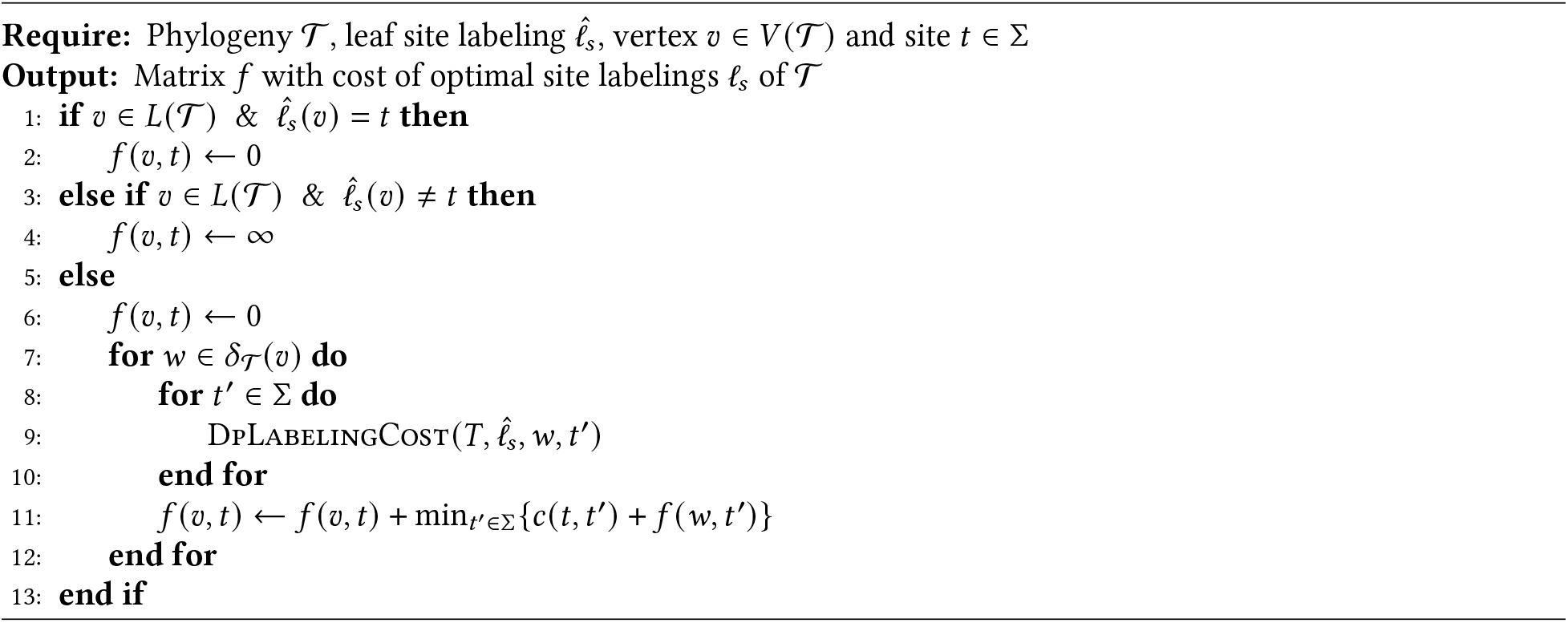

For each vertex 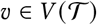 and site *t* ∈ Σ, we define

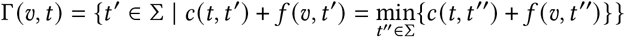

which is the set of labelings for vertex *υ* that are feasible under maximum parsimony when the parent 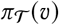 is labeled by *t*. In general, there can be multiple labelings *ℓ_s_* that yield the minimum number of migrations. In order to minimize the number of cross-metastasis seeding events, we find a maximum parsimony site labeling *ℓ_s_* that maximizes the number of nodes labeled by the primary tumor with a single top-down traversal using Algorithm 5.

#### Algorithm 5 DpGetLabeling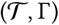

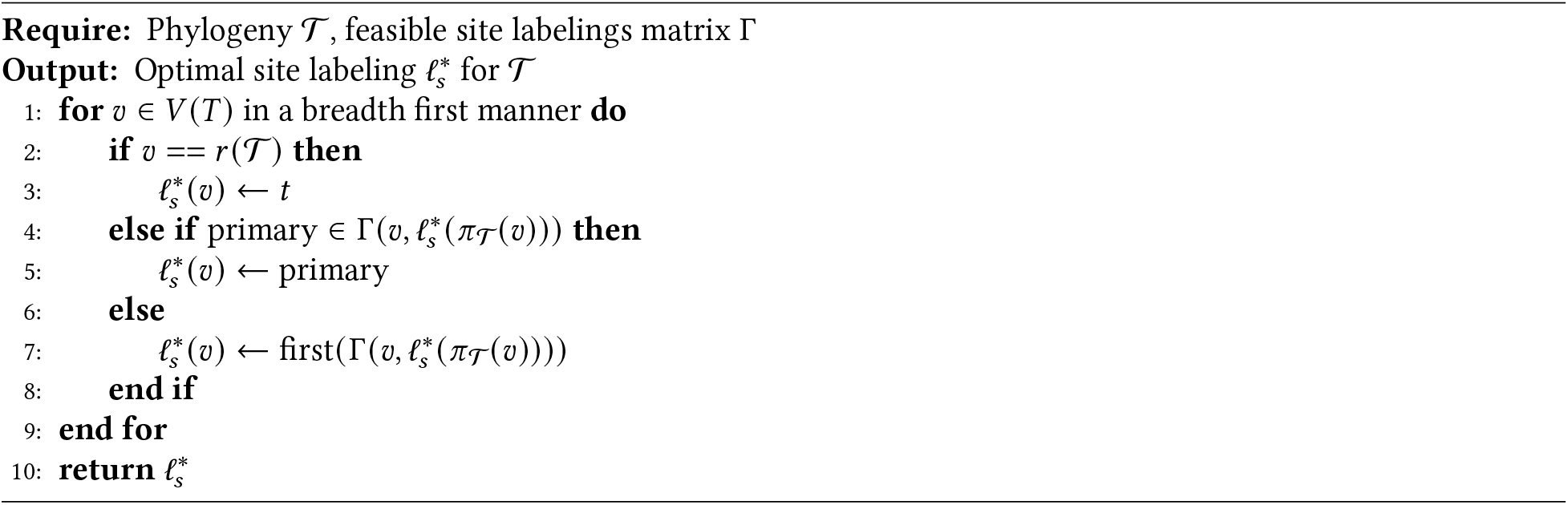

## B Supplementary Results

### B.1 Properties of *k*-star homoplasy phylogenies

We restate and prove a characterization of *k*-star homoplasy phylogenies that is used for constructing our ILP.

#### Theorem 6.

*Character matrix A* ∈ {−1, 0,…, *r*}^*n*×*m*^ *admits a k-star homoplasy phylogeny if and only if there exists a k -star binarization B* ∈ {0, 1}^*n*×*mrk*^ *of A that admits a binary perfect phylogeny*.

*Proof*. (⟹) Suppose *B* is a *k*-star binarization of *A* that admits a binary perfect phylogeny 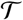. We will then construct a *k*-star homoplasy phylogeny 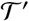 realizing *A* as follows. For each edge 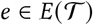 labeled by a mutation (*j, s, p*) → 1, we replace the label of the mutation with *j* → *s* to obtain a phylogeny 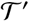. Since 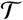 is a perfect phylogeny and *p* ∈ [*k*], each mutation *j* → *s* in 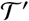 occurs at most *k* times. Further, since in the binarization the 1-sets of (*j, s, p*) and (*j, s, p′*) are disjoint, no mutations (*j, s, p*) and (*j, s, p′*) such that *p* ≠ *p*′ ∈ [*k*] occur along the same path from root in 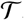. Thus, each mutation (*j, s*) occurs at most once on each path from root in 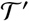; as well. This shows that 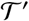 is a *k*-star homoplasy tree. Next, we will show that 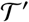 realizes *A*. If *A_i, j_* = *s* ≠ 0 then *B*_*i*, (*j,s,p*_) = 1 for some *p* ∈ [*k*]. Thus, in 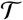 there is some mutation (*j, s, p*) → 1 on the path to cell *i* and therefore a mutation *j* → *s* on the path to cell *i* in 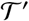. Similarly, if *A_i,j_* = 0 there is no such mutation in 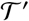. Each path to every *i* is then labeled by all its mutations, showing that 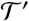 realizes *A*.

(⇐) Suppose we have a *k*-star homoplasy phylogeny 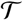 for the matrix *A*. We construct a binary perfect phylogeny 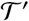 whose character matrix is a binarization of *A* as follows. For each of the at-most *k* edges in 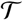 with labels of the form *j* → *s*, we arbitrarily relabel each mutation as (*j, s, p*) → 1 using unique *p* ∈ [*k*] for each edge. Since each mutation (*j, s, p*) occurs at most once in 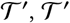 is clearly a binary perfect phylogeny. Let *B* be the unique binary character matrix of 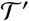. By our construction 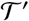 and the star homoplasy property of 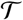, each cell *i* is labeled by exactly one mutation (*j, s, p*) in 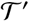 if *A_i,j_* = *s* and no mutations of the form (*j, s, p*) otherwise. Thus *B* is a *k*-star binarization of *A* that admits a binary perfect phylogeny.

The above theorem gives us the following corollary, which is an extension of the three-gamete condition to star perfect phylogeny.

#### Corollary 3.

*Character matrix A admits a star perfect phylogeny if and only if for all pairs of states* (*s, s′*) *such that s,s*′ ∈ {1,…, *r*}, *A does not contain submatrices of the form*

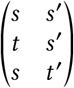

*where t, t*′ ∈{0,…, *r*} *states such that s* ≠ *t and s*′ ≠ *t*′ *for any permutation of rows and columns*.

### B.2 NP-hardness of maximum parsimony under *k*-star homoplasy

We prove the NP-hardness of *k*-star homoplasy for *k* ≥ 4 by a reduction from Cubic Vertex Cover [GP95;JG79]. A cubic graph is a graph in which all vertices have degree 3 (also known as 3-regular graph or trivalent graph). For completeness, we include a description of the problem.

#### Cubic Vertex Cover Problem

**Instance.** A cubic graph *G* = (*V, E*) and a positive integer *k*.
**Question.** Does there exist a vertex cover of *G* of size at most *k*?

Since the problem described in the main text is an optimization problem, we first have to frame it as decision problem to state any type of hardness result. We do this in the obvious way:

#### *k*-star homoplasy problem

**Instance.** A set of *n* species *s_i_*, ∈ {0, 1} and a positive integer *k*.
**Question.** Does there exist a *k*-star homoplasy phylogeny for species 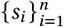 using at most *k* mutations?

We then apply a nearly identical strategy to the reduction from Vertex Cover to binary Camin-Sokal used in [DJS86] to obtain the following theorem.

#### Theorem 7.

*The parsimonious k-star homoplasy decision problem is NP-hard for k* ≥ 4.

*Proof*. Let *G* = ([*m*], *E*) and *k* be an instance of cubic vertex cover. We define the species set *s_e_* ∈ {0, 1}^*m*^ as

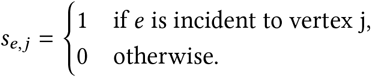

We now claim that there exists a *k*-star homoplasy phylogeny for species set {*s_e_*}_*e*∈E_ using at most *k*′ = |*E*| + *k* mutations if and only if there exists a vertex cover of size at most *k* for *G*.

(⟹) Suppose we have a *k*-star homoplasy phylogeny 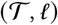 for {*s_e_*}_*e*∈*E*_ using at most *k*′ mutations. Without loss of generality we can assume that 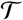 has exactly one mutation on each edge. We define the rank of a vertex 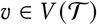 to be the number of 1 entries in its labeling *ℓ*. Since the *k*-homoplasy model is non-modifiable and each edge has exactly one mutation, the rank of each vertex corresponds to its depth in the tree. Since there are |*E*| distinct species of rank-2, there must be |*E*| edges from rank-1 to rank-2 vertices. We associate each rank-1 vertex 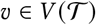 with the unique vertex *i* ∈ [*m*] such that *ℓ*(*υ, i*) = 1. This set of vertices then defines a vertex cover, since every edge *e* = {*i, i*′} corresponds to a species *s_e_* that is a child of either the vertex in 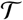 associated with *i* or *i*′. Since, there are at most *m* edges from rank-1 to rank-2 vertices, there are at most *k* rank-1 vertices. Thus, the aforementioned vertex cover has size at most *k*.

(**←**) Suppose we have a vertex cover *S* ⊆ [*m*] of size at most *k*. We define the following phylogenetic tree 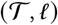 by a vertex set *V*′ = {0} ∪ *S* ∪ *E* and edge set *E*′ that connects the root to every vertex in *S* and each vertex in *e* ∈ *E* to a single vertex *s* ∈ *S* such that *e* is incident to s in *G*. We label the vertices *s* ∈ *S* by the labeling 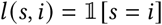 and 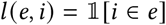. Clearly, the number of mutations from the root to rank-1 vertices is equal to |*S*|. The number of mutations from rank-1 vertices to rank-2 vertices is equal to |*E*| since every vertex *e* ∈ *E* is connected to a vertex *s* ∈ *S* whose labeling differs at exactly one character. The total number of mutations is then at most *k*′ and since it realizes all vertices {*s_e_*}_*e*∈*E*_ it suffices to show that this is a *k*-star homoplasy tree. This last claim follows because the graph is cubic and each vertex has degree 3. That is, a character mutates at most 4 times: once to a rank-1 vertex and three times to rank-2 vertices. Since *k* ≥ 4, this is a *k*-star homoplasy tree.

### B.3 Linear time algorithm for star perfect phylogeny

Theorem 3 tells us that after binary factorization, star perfect phylogeny reduces to binary perfect phylogeny. This hints at a close correspondence between the two problems and it immediately yields a 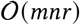 algorithm for star perfect phylogeny. However, the correspondence between the two problems goes deeper. In fact, we can generalize Gusfield’s algorithm to construct a linear time algorithm for star perfect phylogeny. And as part of the proof, we show that there is an analogy of Gusfield’s “Shared Prefix Property” for star perfect phylogeny [Gus14].

#### Theorem 8.

*Algorithm 6 outputs a star perfect phylogeny for character matrix A* ∈ {0,…, *r*}^*n*×*m*^ *in* 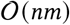 *time if one exists and otherwise outputs HALT*.

*Proof*. We first note that for any species *i* in row *i* of *A*, the path from root to the species is determined up to a permutation of its vertices. Specifically, the path from root to *i* has length *l* where *l* is the number of non-zeros in the row and contains all mutations *j* → *s* in the row of *i*. This is because mutations can only ever be gained and each mutation must occur at least once. It then suffices to find the correct ordering of mutations and then glue the paths of each row together.

To find the correct ordering of a path, take each mutation *m* = (*j, s*) and count the number of times it appears in *A*.Sort these counts and order the mutations by them such that *m*_1_ ≥ *m*_2_ ≥ … ≥ *m_p_*. This should take linear time using radix sort since there are at most *O*(*mn*) mutations and each mutation occurs at most *n* times. Then the order of any given path from root to leaf *i* is determined by this ordering. This is because if mutation *m*_*k*_1__ ≥ *m*_*k*_2__, *m*_*k*_1__ must occur first on any path from the root, otherwise *mk_2_* would have a greater count as mutations only occur once and cannot be reversed.

Now we know that in linear time, we can construct the paths from root to each leaf *i*. It suffices to glue them together to construct our tree. We now argue that Algorithm 6 does this using induction. Namely, we suppose that at the beginning of iteration *i* + 1 of the for-loop, *T_i_* is a star perfect phylogeny for the first *i* rows of *A*.

Suppose that our tree 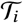 has been constructed for the first *i* rows. From the preceding argument, we know that *p* = (*m*_*σ*(_1_),…, *m*_*σ*(*k*)__) is the path from root down to row *i* + 1. We then follow this path *p* from the root, until the node we are on has no out-edges matching the path. At this point, we glue on the remainder of our path to the node. This will construct a star perfect phylogeny for the first *i* + 1 rows as long as the first *i* + 1 rows of the matrix admit a star perfect phylogeny. If the first *i* + 1 rows do not admit a star perfect phylogeny, it is easy to check by maintaining a set of the added mutations and verifying that any mutation we add does not belong to the set. Thus, Algorithm 6 is a linear time algorithm for star perfect phylogeny.

#### Algorithm 6 Fast star perfect phylogeny algorithm

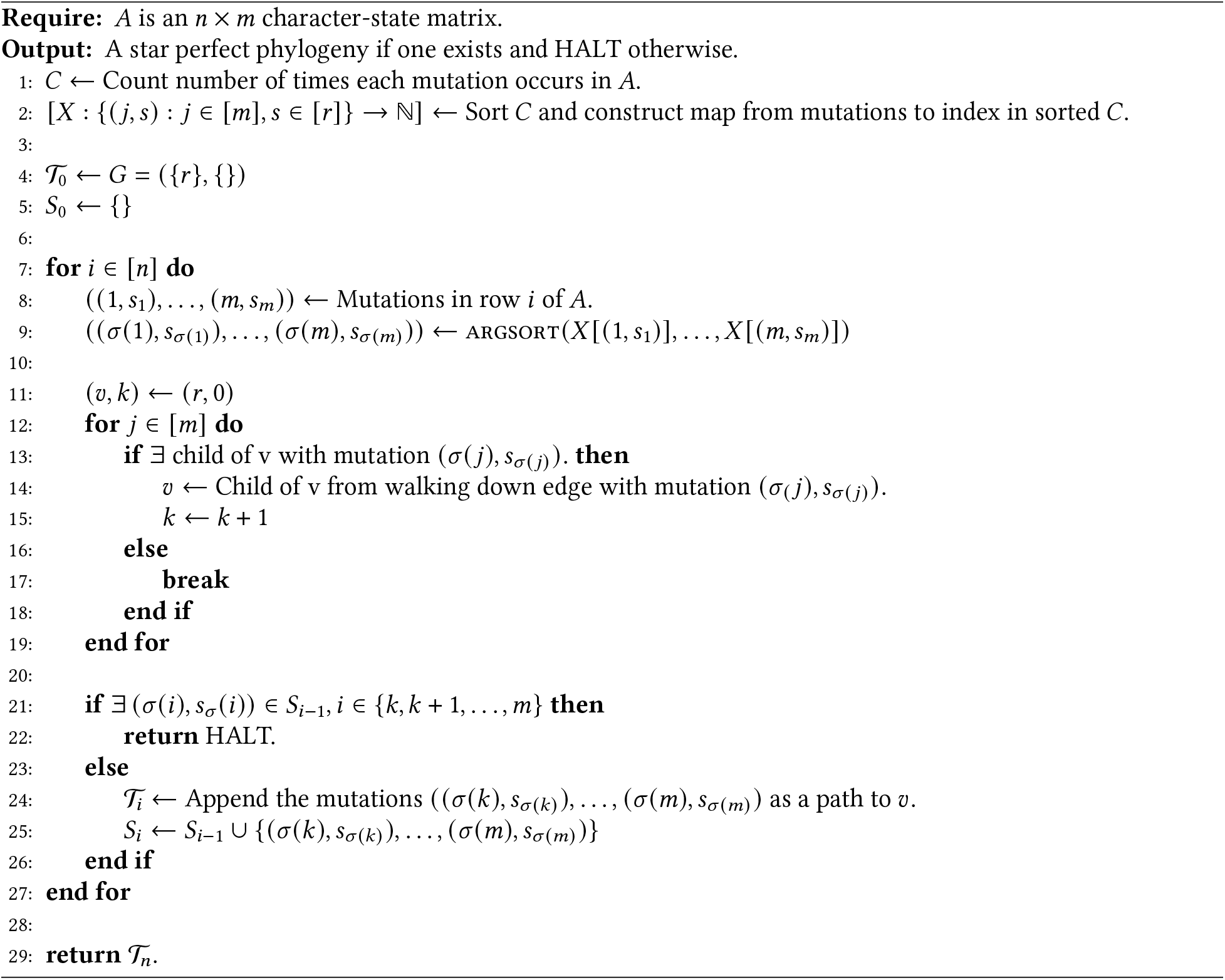

## C Supplementary Figures and Tables

We have the following supplementary figures and tables:

- Table S1 shows time taken by *Startle*, Cassiopeia-ILP and Cassiopeia-Hybrid on medium sized simulated instances (number of characters *m* = 30).
- Table S2 shows the correlation between the RF distance and the star homoplasy parsimony score on simulated instances.
- Supplementary Figure S1 shows the state-transition graphs for five evolutionary models.
- Supplementary Figure S2 compares the performance of *Startle* and existing methods on small instances with no dropout.
- Supplementary Figure S3 compares the performance of *Startle* and existing methods on medium sized instances with varying amount of dropout using RF, Quartet, and Triplet distance metrics.
- Supplementary Figure S4 compares the performance of *Startle* and existing methods on medium sized instances with varying amounts of dropout using RF and weighted star homoplasy parsimony scores.
- Supplementary Figure S5 shows the published phylogeny and *Startle* phylogeny for sample `3513_NT_T1_Fam` with the corresponding migration graphs.

---

**Table S1:** Timing result comparison for inferring medium sized trees with high amounts of dropout. Each instance was evaluated and timed on a 32-core node with an 8-hour time limit as part of the Princeton University Computer Science Cluster Computing Infrastructure. Since the computing cluster is heterogenous, the exact specifications of individual nodes may vary from instance to instance.

| Cells | Characters | Dropout | Seed | Mutation Rate | *Startle*-ILP | Cassiopeia-ILP | Cassiopeia-Hybrid |
| --- | --- | --- | --- | --- | --- | --- | --- |
| 100 | 30 | 0.15 | 10 | 0.2 | **192s** | ≥ 28800s | 12806s |
| 100 | 30 | 0.15 | 11 | 0.2 | **214s** | ≥ 28800s | 13231s |
| 100 | 30 | 0.15 | 12 | 0.2 | **119s** | ≥ 28800s | 10389s |
| 100 | 30 | 0.2 | 10 | 0.2 | **243s** | ≥ 28800s | ≥ 28800s |
| 100 | 30 | 0.2 | 11 | 0.2 | **394s** | ≥ 28800s | ≥ 28800s |
| 100 | 30 | 0.2 | 12 | 0.2 | **184s** | ≥ 28800s | ≥ 28800s |
| 150 | 30 | 0.15 | 10 | 0.2 | 382s | ≥ 28800s | **107s** |
| 150 | 30 | 0.15 | 11 | 0.2 | **187s** | ≥ 28800s | 12858 |
| 150 | 30 | 0.15 | 12 | 0.2 | **395s** | ≥ 28800s | 4845 |
| 150 | 30 | 0.2 | 10 | 0.2 | **471s** | ≥ 28800s | ≥ 28800s |
| 150 | 30 | 0.2 | 11 | 0.2 | **279s** | ≥ 28800s | ≥ 28800s |
| 150 | 30 | 0.2 | 12 | 0.2 | **509s** | ≥ 28800s | ≥ 28800s |
| 200 | 30 | 0.15 | 10 | 0.2 | 651s | 24339s | **68s** |
| 200 | 30 | 0.15 | 11 | 0.2 | **2072s** | ≥ 28800s | 25675s |
| 200 | 30 | 0.15 | 12 | 0.2 | **686s** | 15732s | 12325s |
| 200 | 30 | 0.2 | 10 | 0.2 | **1455s** | ≥ 28800s | ≥ 28800s |
| 200 | 30 | 0.2 | 11 | 0.2 | **1978s** | ≥ 28800s | ≥ 28800s |
| 200 | 30 | 0.2 | 12 | 0.2 | **922s** | ≥ 28800s | ≥ 28800s |
| 50 | 30 | 0.15 | 10 | 0.2 | **111s** | 5641s | 12682s |
| 50 | 30 | 0.15 | 11 | 0.2 | **92s** | 8360s | 11818s |

**Table S2:** Rank correlation between median RF distance and median star homoplasy Weighted Parsimony scores. Specifically, we computed the rank of each method for a specific parameter setting using both the median RF distance and median Parsimony criterion and then computed the PearsonR correlation between the two lists of ranks.

| Cells | Characters | Dropout | SpearmanR | p-value |
| --- | --- | --- | --- | --- |
| 50 | 30 | 0.05 | 0.80 | 0.102 |
| 100 | 10 | 0.05 | 0.89 | 0.041 |
| 100 | 20 | 0.05 | 0.79 | 0.104 |
| 100 | 30 | 0.05 | 0.87 | 0.054 |
| 150 | 30 | 0.05 | 0.90 | 0.037 |
| 200 | 30 | 0.05 | 0.10 | 0.8729 |
| 50 | 20 | 0.15 | 0.92 | 0.028 |
| 50 | 30 | 0.15 | 0.67 | 0.219 |
| 100 | 10 | 0.15 | 0.82 | 0.089 |
| 100 | 20 | 0.15 | 0.97 | 0.005 |
| 100 | 30 | 0.15 | 0.80 | 0.200 |
| 150 | 30 | 0.15 | 0.90 | 0.037 |
| 200 | 30 | 0.15 | 0.80 | 0.200 |
| 50 | 20 | 0.20 | 0.92 | 0.028 |
| 50 | 30 | 0.20 | 1.00 | 0.0 |
| 100 | 10 | 0.20 | 0.90 | 0.037 |
| 100 | 20 | 0.20 | 1.00 | 0.0 |
| 100 | 30 | 0.20 | 0.80 | 0.200 |
| 150 | 30 | 0.20 | 1.00 | 0.0 |
| 200 | 30 | 0.20 | 1.00 | 0.0 |

**Supplementary Figure S1:**
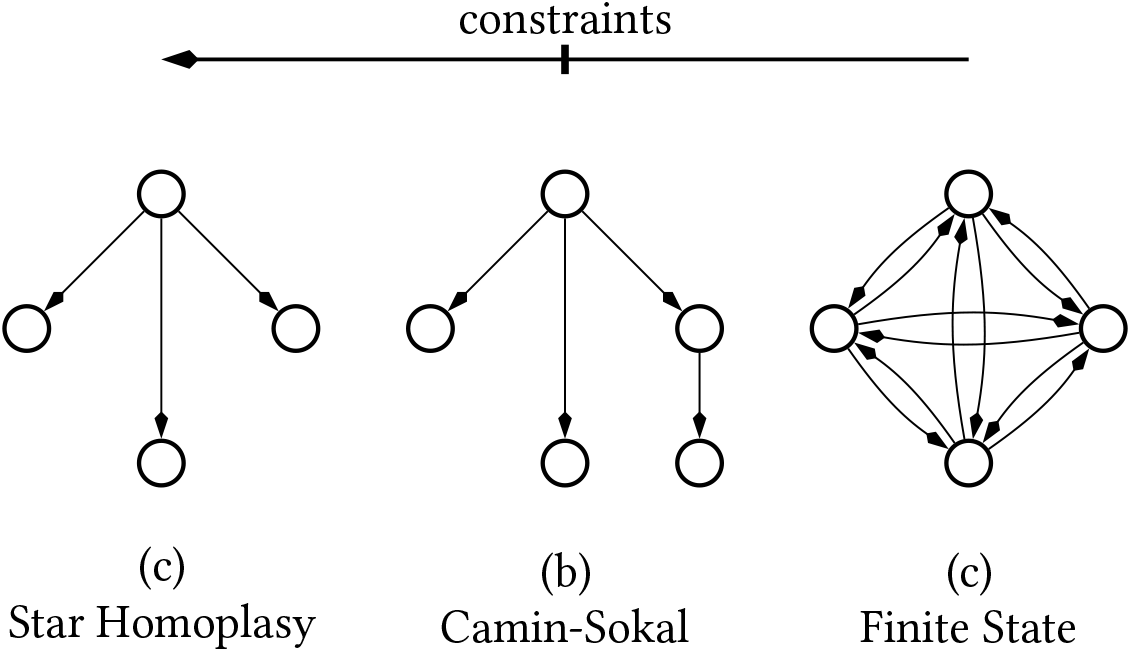
State-transition graphs for three evolutionary models with strictly decreasing evolutionary constraints. Edges denote the allowed set of state transitions.

**Supplementary Figure S2:**
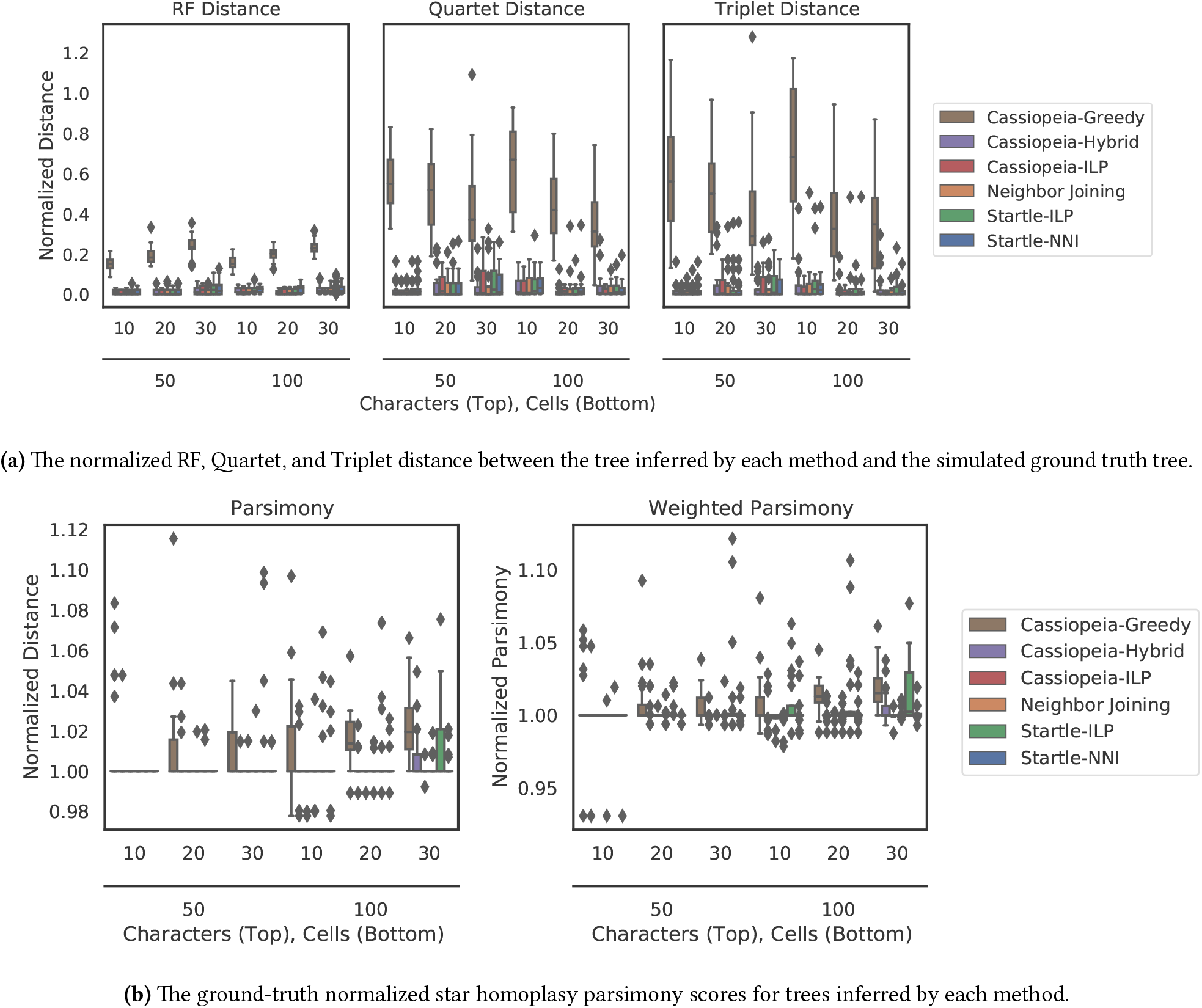
Performance comparison on Cassiopeia simulated data for a small number of cells and characters with no dropout at mutation rate *p* = 0.1.

**Supplementary Figure S3:**
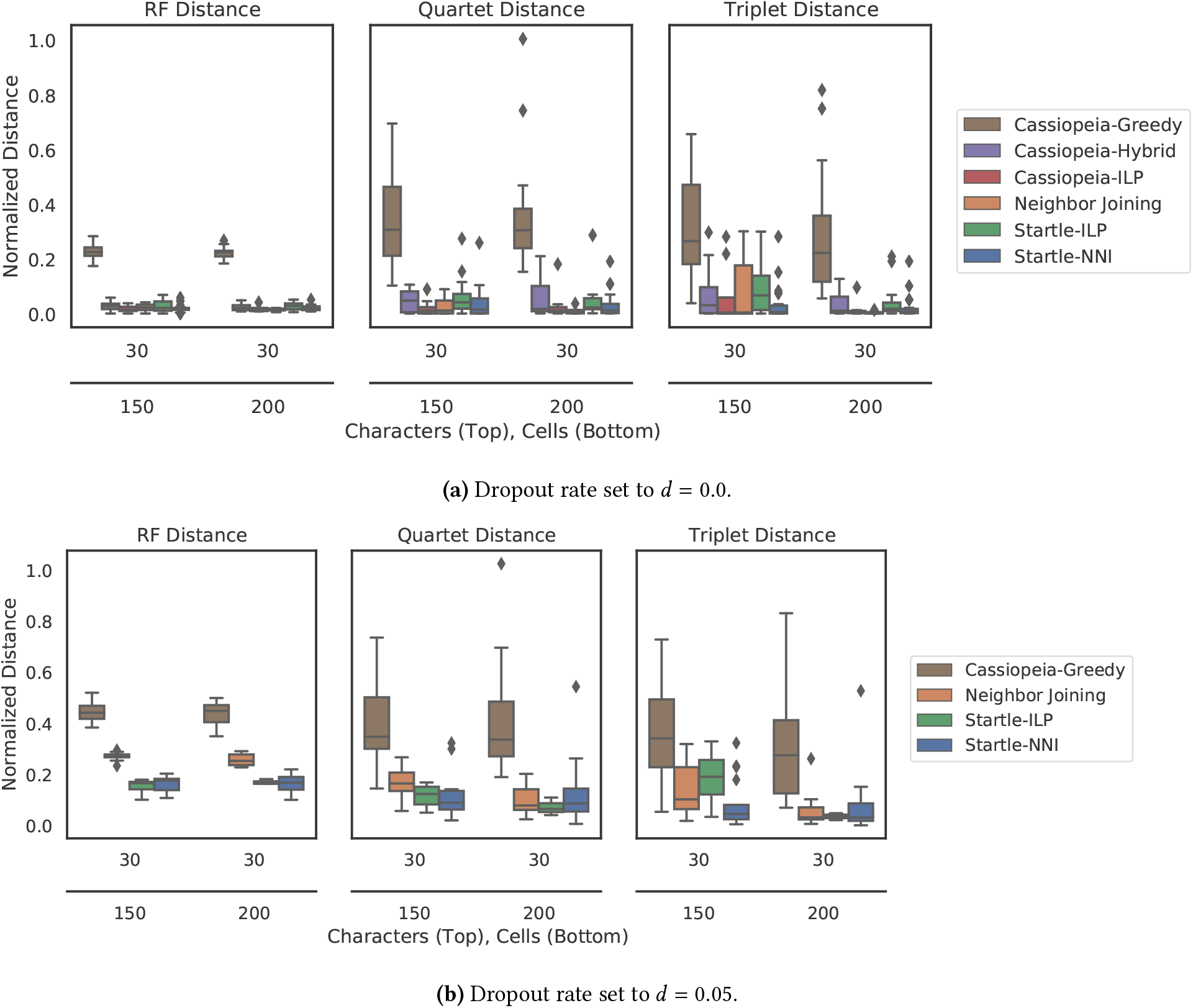
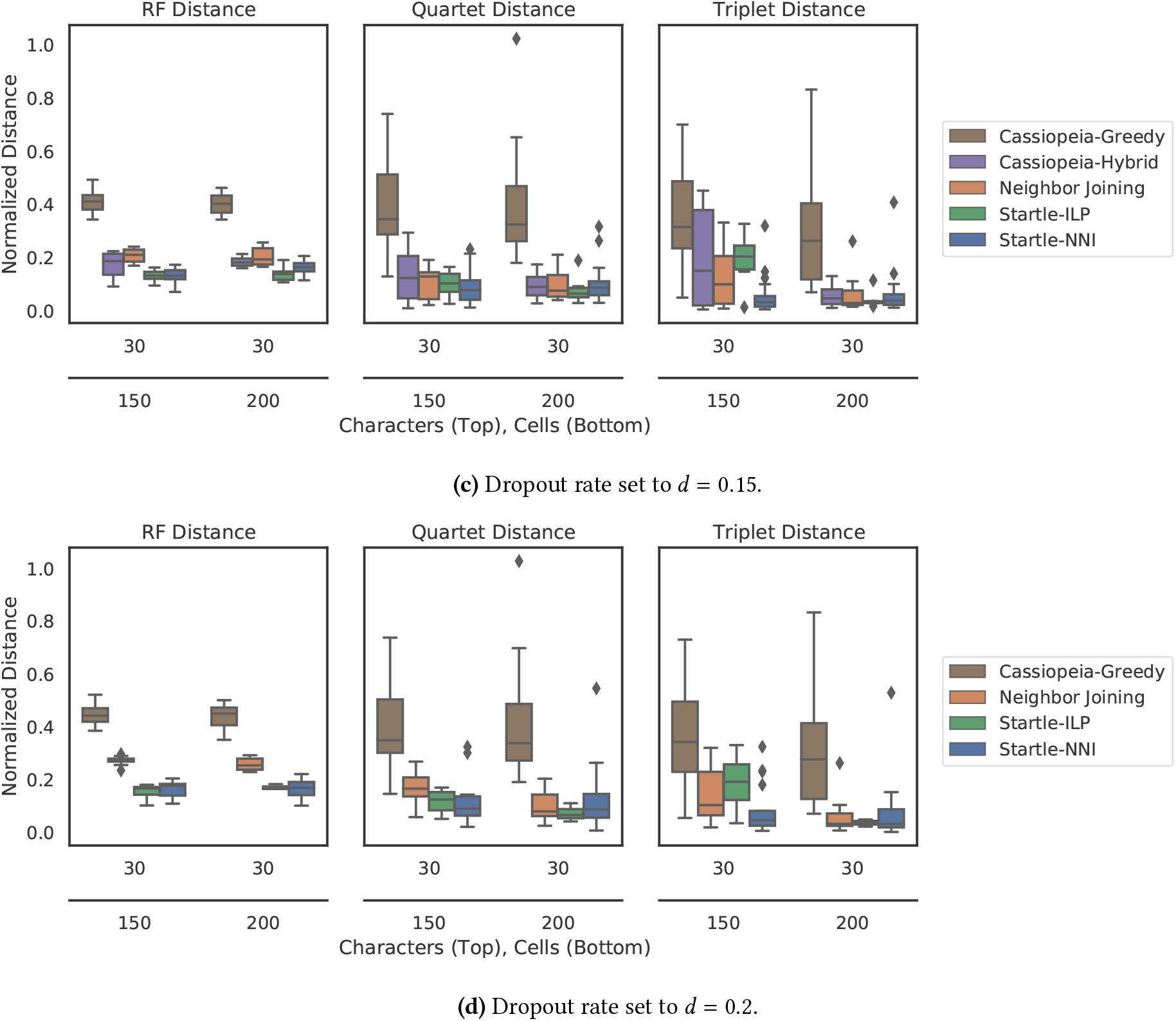
The normalized RF, Quartet, and Triplet distance between the tree inferred by each method and the simulated ground truth tree on medium sized instances with varying amounts of dropout. On instances with dropout *d* ≥ 0.05, Cassiopeia-Hybrid and Cassiopeia-ILP did not terminate in time and are not included.

**Supplementary Figure S4:**
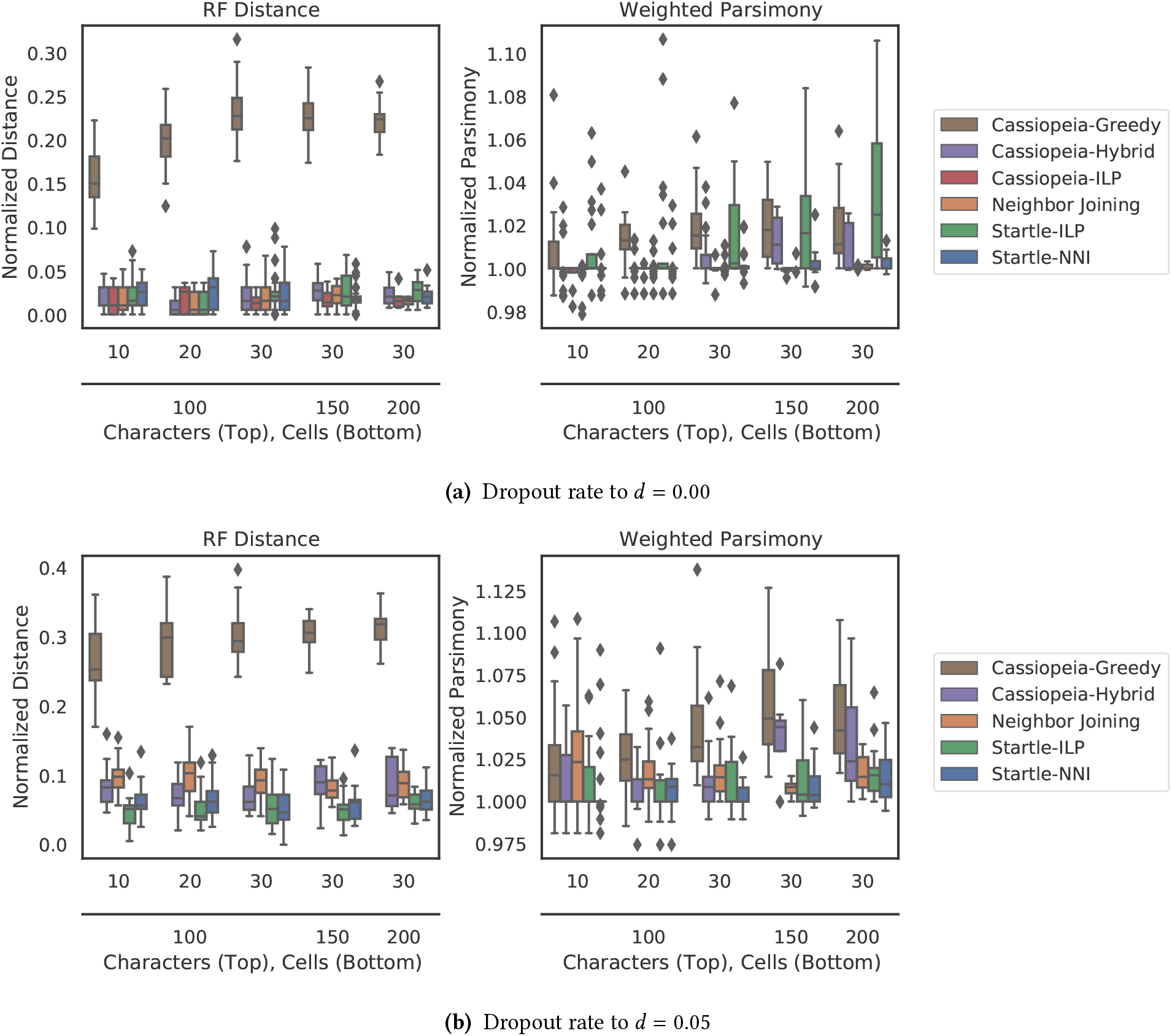
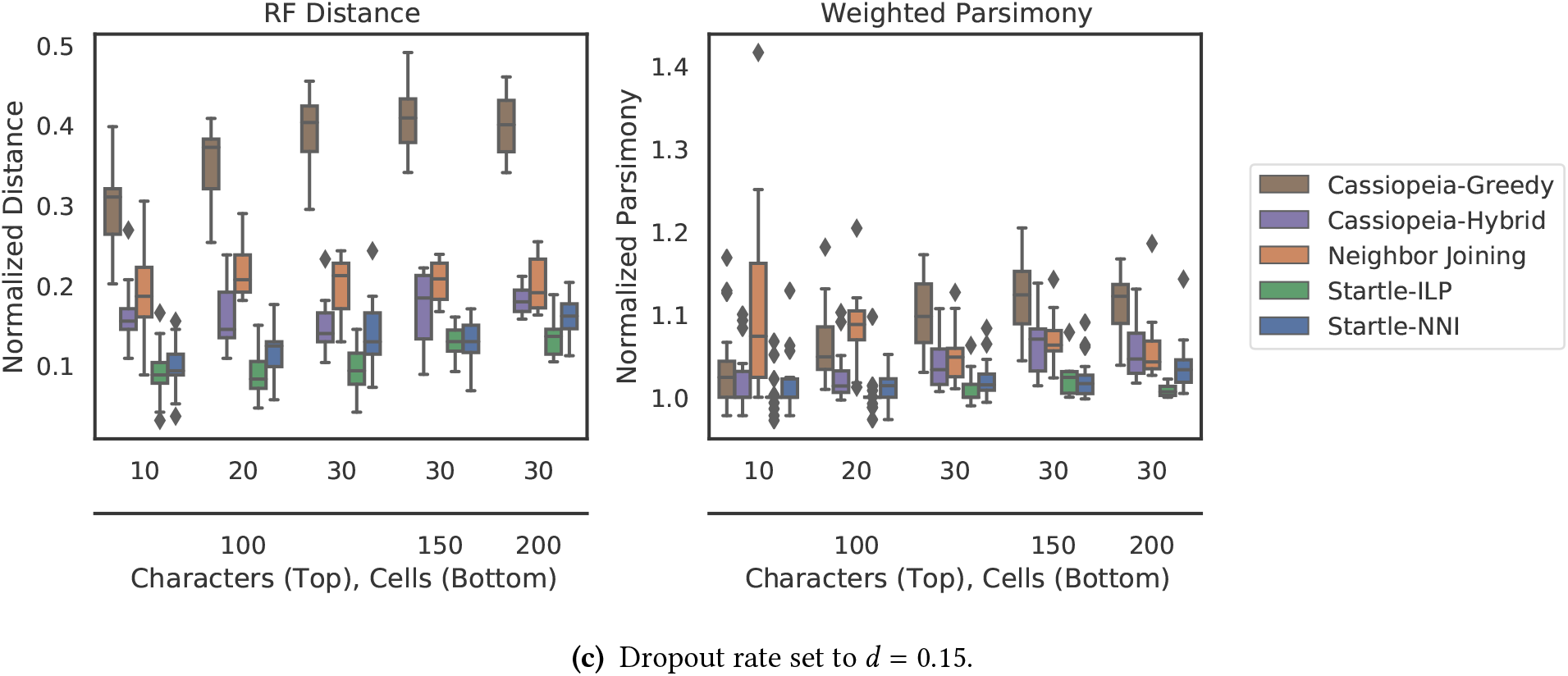
(Left) The normalized Robinson-Foulds (RF) distance between the tree inferred by each method and the simulated ground truth tree. (Right) The weighted star homoplasy parsimony score for each method, normalized such that the simulated tree has score = 1. Simulated datasets were generated with mutation rate *p* = 0.1 and a varying dropout rate *d* ∈ {0.00, 0.05, 0.15}.

**Supplementary Figure S5:**
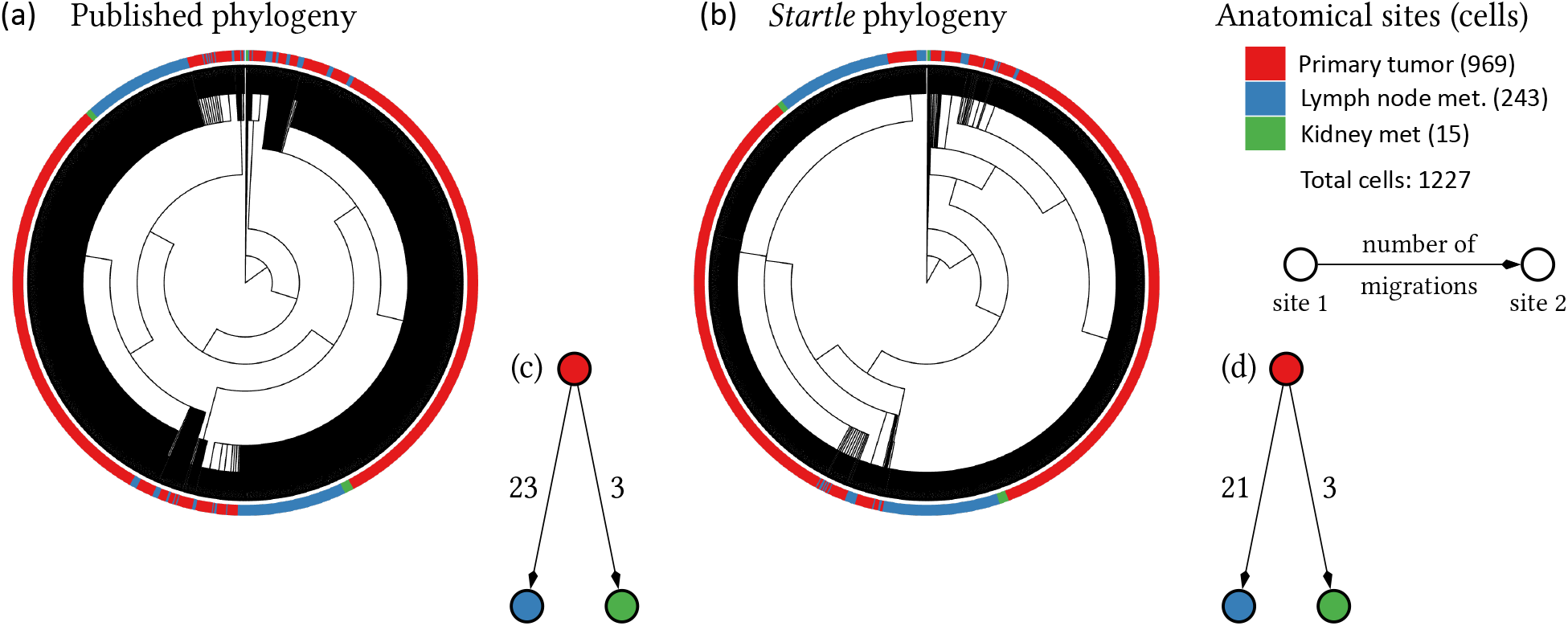
(a) Published phylogeny and (b) *Startle* phylogeny for sample 3513_NT_T1_Fam. (c) MACHINA migration graphs inferred using the published phylogeny and (d) *Startle* phylogeny. Directed edges indicate the direction of migration between anatomical sites and the edge weights denote the number of cells that migrated between the anatomical sites.

## Footnotes

1 These graphs are also sometimes defined with weighted edges, where the weight of an edges indicates the cost of the associated state transition.

2 This assumption is also called the infinite alleles assumption, or the infinitely-many alleles assumption [Fer01;Gus14;Ewe16]

3 In the original study, [Yan+22] analyzed metastasis by computing a normalized measure of the phylogenetic distance between the metastatic cells and the primary tumor cells. However, this approach does not provide insight about the migration history, i.e. the direction of migration of cells and the number of clones that migrated during metastasis.

## Bibliography

[1] Anna Alemany et al. “Whole-organism clone tracing using single-cell sequencing”. In: Nature 556.7699 (2018), pp. 108–112.

[2] Andrew V Anzalone, Luke W Koblan, and David R Liu. “Genome editing with CRISPR–Cas nucleases, base editors, transposases and prime editors”. In: Nature biotechnology 38.7 (2020), pp. 824–844.

[3] Sam Behjati et al. “Genome sequencing of normal cells reveals developmental lineages and mutational processes”. In: Nature 513.7518 (2014), pp. 422–425.

[4] Damian Bogdanowicz and Krzysztof Giaro. “Matching split distance for unrooted binary phylogenetic trees”. In: IEEE/ACM Transactions on Computational Biology and Bioinformatics 9.1 (2011), pp. 150–160.

[5] Damian Bogdanowicz, Krzysztof Giaro, and Borys Wróbel. “TreeCmp: comparison of trees in polynomial time”. In: Evolutionary Bioinformatics 8 (2012), EBO–S9657.

[6] Paola Bonizzoni et al. “Beyond perfect phylogeny: Multisample phylogeny reconstruction via ilp”. In: Proceedings of the 8th ACM International Conference on Bioinformatics, Computational Biology, and Health Informatics. 2017, pp. 1–10.

[7] Yehuda Brody et al. “Quantification of somatic mutation flow across individual cell division events by lineage sequencing”. In: Genome research 28.12 (2018), pp. 1901–1918.

[8] Joseph H Camin and Robert R Sokal. “A method for deducing branching sequences in phylogeny”. In: Evolution (1965), pp. 311–326.

[9] Gabriel Cardona, Francesc Rosselló, and Gabriel Valiente. “Extended Newick: it is time for a standard representation of phylogenetic networks”. In: BMC bioinformatics 9.1 (2008), pp. 1–8.

[10] Cheryl A Carlson et al. “Decoding cell lineage from acquired mutations using arbitrary deep sequencing”. In: Nature methods 9.1 (2012), pp. 78–80.

[11] Michelle M Chan et al. “Molecular recording of mammalian embryogenesis”. In: Nature 570.7759 (2019), pp. 77–82.

[12] Markus Chimani, Sven Rahmann, and Sebastian Böcker. “Exact ILP solutions for phylogenetic minimum flip problems”. In: Proceedings of the First ACM International Conference on Bioinformatics and Computational Biology. 2010, pp. 147–153.

[13] Simone Ciccolella et al. “gpps: an ILP-based approach for inferring cancer progression with mutation losses from single cell data”. In: BMC bioinformatics 21.1 (2020), pp. 1–16.

[14] Douglas E Critchlow, Dennis K Pearl, and Chunlin Qian. “The triples distance for rooted bifurcating phylogenetic trees”. In: Systematic Biology 45.3 (1996), pp. 323–334.

[15] William HE Day, David S Johnson, and David Sankoff. “The computational complexity of inferring rooted phylogenies by parsimony”. In: Mathematical biosciences 81.1 (1986), pp. 33–42.

[16] Amit G Deshwar et al. “PhyloWGS: reconstructing subclonal composition and evolution from whole-genome sequencing of tumors”. In: Genome biology 16.1 (2015), pp. 1–20.

[17] George F Estabrook, FR McMorris, and Christopher A Meacham. “Comparison of undirected phylogenetic trees based on subtrees of four evolutionary units”. In: Systematic Zoology 34.2 (1985), pp. 193–200.

[18] Warren J Ewens. “Motoo Kimura and James Crow on the Infinitely Many Alleles Model”. In: Genetics 202.4 (2016), pp. 1243–1245.

[19] James S Farris. “Methods for computing Wagner trees”. In: Systematic Biology 19.1 (1970), pp. 83–92.

[20] Joseph Felsenstein. “Cases in which parsimony or compatibility methods will be positively misleading”. In: Systematic zoology 27.4 (1978), pp. 401–410.

[21] Joseph Felsenstein. PHYLIP (phylogeny inference package), version 3.5 c. Joseph Felsenstein., 1993.

[22] Joseph Felsenstein and Joseph Felenstein. Inferring phylogenies. Vol. 2. Sinauer associates Sunderland, MA, 2004.

[23] Jean Feng et al. “Estimation of cell lineage trees by maximum-likelihood phylogenetics”. In: The annals of applied statistics 15.1 (2021), p. 343.

[24] David Fernández-Baca. “The perfect phylogeny problem”. In: Steiner Trees in Industry. Springer, 2001, pp. 203–234.

[25] Walter M Fitch. “Toward defining the course of evolution: minimum change for a specific tree topology”. In: Systematic Biology 20.4 (1971), pp. 406–416.

[26] Wuming Gong et al. “Benchmarked approaches for reconstruction of in vitro cell lineages and in silico models of C. elegans and M. musculus developmental trees”. In: Cell systems 12.8 (2021), pp. 810–826.

[27] Wuming Gong et al. “Single cell lineage reconstruction using distance-based algorithms and the R package, DCLEAR”. In: BMC bioinformatics 23.1 (2022), pp. 1–14.

[28] Raymond Greenlaw and Rossella Petreschi. “Cubic graphs”. In: ACM Computing Surveys (CSUR) 27.4 (1995), pp. 471–495.

[29] Stéphane Guindon et al. “New algorithms and methods to estimate maximum-likelihood phylogenies: assessing the performance of PhyML 3.0”. In: Systematic biology 59.3 (2010), pp. 307–321.

[30] Dan Gusfield. “Efficient algorithms for inferring evolutionary trees”. In: Networks 21.1 (1991), pp. 19–28.

[31] Dan Gusfield. ReCombinatorics: the algorithmics of ancestral recombination graphs and explicit phylogenetic networks. MIT press, 2014.

[32] Katharina Jahn, Jack Kuipers, and Niko Beerenwinkel. “Tree inference for single-cell data”. In: Genome biology 17.1 (2016), pp. 1–17.

[33] Yuchao Jiang et al. “Assessing intratumor heterogeneity and tracking longitudinal and spatial clonal evolutionary history by next-generation sequencing”. In: Proceedings of the National Academy of Sciences 113.37 (2016), E5528–E5537.

[34] David S Johnson and Michael R Garey. Computers and intractability: A guide to the theory of NP-completeness. WH Freeman, 1979.

[35] Matthew G Jones et al. “Inference of single-cell phylogenies from lineage tracing data using Cassiopeia”. In: Genome biology 21.1 (2020), pp. 1–27.

[36] Reza Kalhor et al. “Developmental barcoding of whole mouse via homing CRISPR”. In: Science 361.6405 (2018), eaat9804.

[37] Mohammed El-Kebir. “SPhyR: tumor phylogeny estimation from single-cell sequencing data under loss and error”. In: Bioinformatics 34.17 (2018), pp. i671–i679.

[38] Mohammed El-Kebir, Gryte Satas, and Benjamin J Raphael. “Inferring parsimonious migration histories for metastatic cancers”. In: Nature genetics 50.5 (2018), pp. 718–726.

[39] Mohammed El-Kebir et al. “Reconstruction of clonal trees and tumor composition from multi-sample sequencing data”. In: Bioinformatics 31.12 (2015), pp. i62–i70.

[40] Naoki Konno et al. “Deep distributed computing to reconstruct extremely large lineage trees”. In: Nature Biotechnology 40.4 (2022), pp. 566–575.

[41] Michael A Lodato et al. “Somatic mutation in single human neurons tracks developmental and transcriptional history”. In: Science 350.6256 (2015), pp. 94–98.

[42] Salem Malikic et al. “PhISCS: a combinatorial approach for subperfect tumor phylogeny reconstruction via integrative use of single-cell and bulk sequencing data”. In: Genome research 29.11 (2019), pp. 1860–1877.

[43] Aaron McKenna and James A Gagnon. “Recording development with single cell dynamic lineage tracing”. In: Development 146.12 (2019), dev169730.

[44] Aaron McKenna et al. “Whole-organism lineage tracing by combinatorial and cumulative genome editing”. In: Science 353.6298 (2016), aaf7907.

[45] Charles D Michener and Robert R Sokal. “A quantitative approach to a problem in classification”. In: Evolution 11.2 (1957), pp. 130–162.

[46] Lam-Tung Nguyen et al. “IQ-TREE: a fast and effective stochastic algorithm for estimating maximum-likelihood phylogenies”. In: Molecular biology and evolution 32.1 (2015), pp. 268–274.

[47] Itsik Pe’er, Ron Shamir, and Roded Sharan. “Incomplete directed perfect phylogeny”. In: Annual Symposium on Combinatorial Pattern Matching. Springer. 2000, pp. 143–153.

[48] Samuel D Perli, Cheryl H Cui, and Timothy K Lu. “Continuous genetic recording with self-targeting CRISPR-Cas in human cells”. In: Science 353.6304 (2016), aag0511.

[49] Victoria Popic et al. “Fast and scalable inference of multi-sample cancer lineages”. In: Genome biology 16.1 (2015), pp. 1–17.

[50] Bushra Raj, James A Gagnon, and Alexander F Schier. “Large-scale reconstruction of cell lineages using singlecell readout of transcriptomes and CRISPR–Cas9 barcodes by scGESTALT”. In: Nature protocols 13.11 (2018), pp. 2685–2713.

[51] Bushra Raj et al. “Simultaneous single-cell profiling of lineages and cell types in the vertebrate brain”. In: Nature biotechnology 36.5 (2018), pp. 442–450.

[52] David F Robinson and Leslie R Foulds. “Comparison of phylogenetic trees”. In: Mathematical biosciences 53.1-2 (1981), pp. 131–147.

[53] Stuart J Russell. Artificial intelligence a modern approach. Pearson Education, Inc., 2010.

[54] Naruya Saitou and Masatoshi Nei. “The neighbor-joining method: a new method for reconstructing phylogenetic trees” In: Molecular biology and evolution 4.4 (1987), pp. 406–425.

[55] David Sankoff and Pascale Rousseau. “Locating the vertices of a Steiner tree in an arbitrary metric space”. In: Mathematical Programming 9.1 (1975), pp. 240–246.

[56] Gryte Satas et al. “SCARLET: single-cell tumor phylogeny inference with copy-number constrained mutation losses”. In: Cell Systems 10.4 (2020), pp. 323–332.

[57] Bastiaan Spanjaard et al. “Simultaneous lineage tracing and cell-type identification using CRISPR–Cas9-induced genetic scars”. In: Nature biotechnology 36.5 (2018), pp. 469–473.

[58] Alexandros Stamatakis. “RAxML version 8: a tool for phylogenetic analysis and post-analysis of large phylogenies”. In: Bioinformatics 30.9 (2014), pp. 1312–1313.

[59] John E Sulston et al. “The embryonic cell lineage of the nematode Caenorhabditis elegans”. In: Developmental biology 100.1 (1983), pp. 64–119.

[60] David L Swofford and Wayne P Maddison. “Parsimony, character-state reconstructions, and evolutionary inferences”. In: Systematics, historical ecology, and North American freshwater fishes 1 (1992).

[61] Liming Tao et al. “Retrospective cell lineage reconstruction in humans by using short tandem repeats”. In: Cell reports methods 1.3 (2021), p. 100054.

[62] Daniel E Wagner and Allon M Klein. “Lineage tracing meets single-cell omics: opportunities and challenges”. In: Nature Reviews Genetics 21.7 (2020), pp. 410–427.

[63] Daniel E Wagner et al. “Single-cell mapping of gene expression landscapes and lineage in the zebrafish embryo”. In: Science 360.6392 (2018), pp. 981–987.

[64] Dian Yang et al. “Lineage tracing reveals the phylodynamics, plasticity, and paths of tumor evolution”. In: Cell 185.11 (2022), pp. 1905–1923.

[65] Hamim Zafar, Chieh Lin, and Ziv Bar-Joseph. “Single-cell lineage tracing by integrating CRISPR-Cas9 mutations with transcriptomic data”. In: Nature communications 11.1 (2020), pp. 1–14.

[66] Leonid Zosin and Samir Khuller. “On directed Steiner trees”. In: SODA. Vol. 2. Citeseer. 2002, pp. 59–63.

